# Seizure pathways change on circadian and slower timescales in individual patients with focal epilepsy

**DOI:** 10.1101/661371

**Authors:** Gabrielle M Schroeder, Beate Diehl, Fahmida A Chowdhury, John S Duncan, Jane de Tisi, Andrew J Trevelyan, Rob Forsyth, Andrew Jackson, Peter N Taylor, Yujiang Wang

## Abstract

Personalised medicine requires that treatments adapt to not only the patient, but changing factors within each individual. Although epilepsy is a dynamic disorder that is characterised by pathological fluctuations in brain state, surprisingly little is known about whether and how seizures vary in the same patient. We quantitatively compared within-patient seizure network dynamics using intracranial recordings of over 500 seizures from 31 patients with focal epilepsy (mean 16.5 seizures/patient). In all patients, we found variability in seizure paths through the space of possible network dynamics, producing either a spectrum or clusters of different dynamics. Seizures with similar pathways tended to occur closer together in time, and a simple model suggested that seizure pathways change on circadian and/or slower timescales in the majority of patients. These temporal relationships occurred independent of whether the patient underwent antiepileptic medication reduction. Our results suggest that various modulatory processes, operating at different timescales, shape within-patient seizure dynamics, leading to variable seizure pathways that may require tailored treatment approaches.

## Introduction

Focal epilepsy is characterised by spontaneous, recurrent seizures that arise from localised cortical sites (1). An unresolved question is how much seizure dynamics can vary in individual patients. Past studies suggest that seizures within a single patient share common features (2–6) and progress through a similar sequence (7), or “characteristic pathway” (8), of neural dynamics. However, there is also evidence that seizure dynamics vary in some patients. Clinically, there may be different types of seizure dynamics in patients with multiple seizure onset sites (9), and long-term electroencephalographic (EEG) recordings suggest that a subset of patients have multiple seizure populations with distinct dynamics (8, 10–12). Ictal onset patterns (13, 14), the extent of seizure spread (15, 16), and seizure recruitment patterns (17) can also differ in the same patient. This variability may arise from fluctuations in the underlying brain state (18–22), suggesting that background neural dynamics affect not only seizure likelihood (19, 23), but also seizure *features*. Crucially, a given treatment may only address a subset of a patient’s seizure dynamics: for example, a single neurostimulation protocol may not control the complete repertoire of seizures (18) and a single prediction algorithm may fail to forecast all seizures (10, 12, 24). Consequently, seizure variability has important implications for clinical management in these patients.

To design optimal and comprehensive treatments, we therefore need to understand the prevalence and characteristics of within-patient seizure variability. Is seizure variability present in all patients, and, if so, what form does the variability take? Do within-patient seizures cluster into groups with distinct dynamics? How are different seizure dynamics distributed in time?

To answer these questions, we must objectively quantify seizure similarity. This task is challenging due to the complexity of seizure dynamics: a variety of spatiotemporal features change independently during seizure evolution. Although some studies have quantitatively compared within-patient seizures (25–30), the current gold standard remains visual inspection of ictal EEG by trained clinicians. This latter approach is time-consuming and subjective, and can miss important features, including functional network interactions, that are difficult to detect visually. These functional network dynamics, also known as functional connectivity patterns, describe relationships between the activity recorded by different EEG channels. Temporal changes in network dynamics play important roles in seizure initiation, propagation, and termination (2, 22, 31–40), in part due to dynamic changes in the connectivity of the seizure onset zone (7, 41–43). To fully understand how functional interactions support ictal processes, we must also determine if multiple seizure pathways, representing different ictal network evolutions, can co-exist in an individual patient. Such diversity would reveal that the same neural regions can variably interact to produce a variety of pathological dynamics.

Our goal was to quantify and characterise within-patient variability in seizure pathways through network space. We visualised and compared the within-patient seizure network evolutions of human patients with focal epilepsy (recorded for 43-382 hrs). In total, we analysed the network evolutions of 511 seizures (average 16.5 seizures/patient), making our study the first large-scale examination of within-patient seizure variability. In each patient, we found variability in seizure network evolution, revealing that within-patient seizures are not well-represented by a single characteristic pathway. However, seizures can share parts or all of the same pathway, with recurring dynamical elements across seizures. Furthermore, we explored how seizure dynamics change over different timescales, providing novel insight into the temporal changes of within-patient seizures. Our analysis revealed that seizures change on circadian and/or slower timescales in each patient, suggesting that different modulatory processes shape seizure pathways.

## Results

We analysed seizure network evolution in 31 human patients (511 seizures total, mean 16.5 seizures/patient) with focal epilepsy who underwent continuous intracranial electroencephalographic (iEEG) recordings as part of presurgical evaluation. Patient details are provided in SI Appendix, Text S1. We first visualise seizure network dynamics and quantify the dissimilarity of within-subject seizure pathways through network space. Importantly, our analysis captures differences in network interactions during seizures, which do not necessarily correspond to anatomical differences in the location and spread of seizure activity. We then investigate the amount and form of this variability across patients, and explore how seizure dynamics change over time. Finally, we hypothesise how underlying processes occurring on different time scales could drive the observed changes in seizure pathways.

### Visualising and quantifying variability in within-patient seizure pathways

Our first goal was to objectively compare within-patient seizure network dynamics. For each patient, we extracted the seizure iEEGs (Fig. 1A) and computed the sliding-window functional connectivity, defined as band-averaged coherence in six frequency bands (Fig. 1B). Thus, each seizure time window was described by a set of six connectivity matrices that captured interactions between iEEG channels in each frequency band. We additionally normalised the magnitude of each connectivity matrix to focus on the evolving patterns of network interactions, rather than gross changes in the global level of coherence. The set of all possible connectivity patterns created a high-dimensional space, in which each location corresponded to a specific network configuration. As such, each time window could be represented by a high-dimensional data point, and the evolution of a seizure’s network dynamics formed a pathway in this high-dimensional connectivity space. By transforming seizures in this manner, we framed our comparison of seizure dynamics as a comparison of seizure pathways (or trajectories) through the high-dimensional network space.

**Fig. 1:**
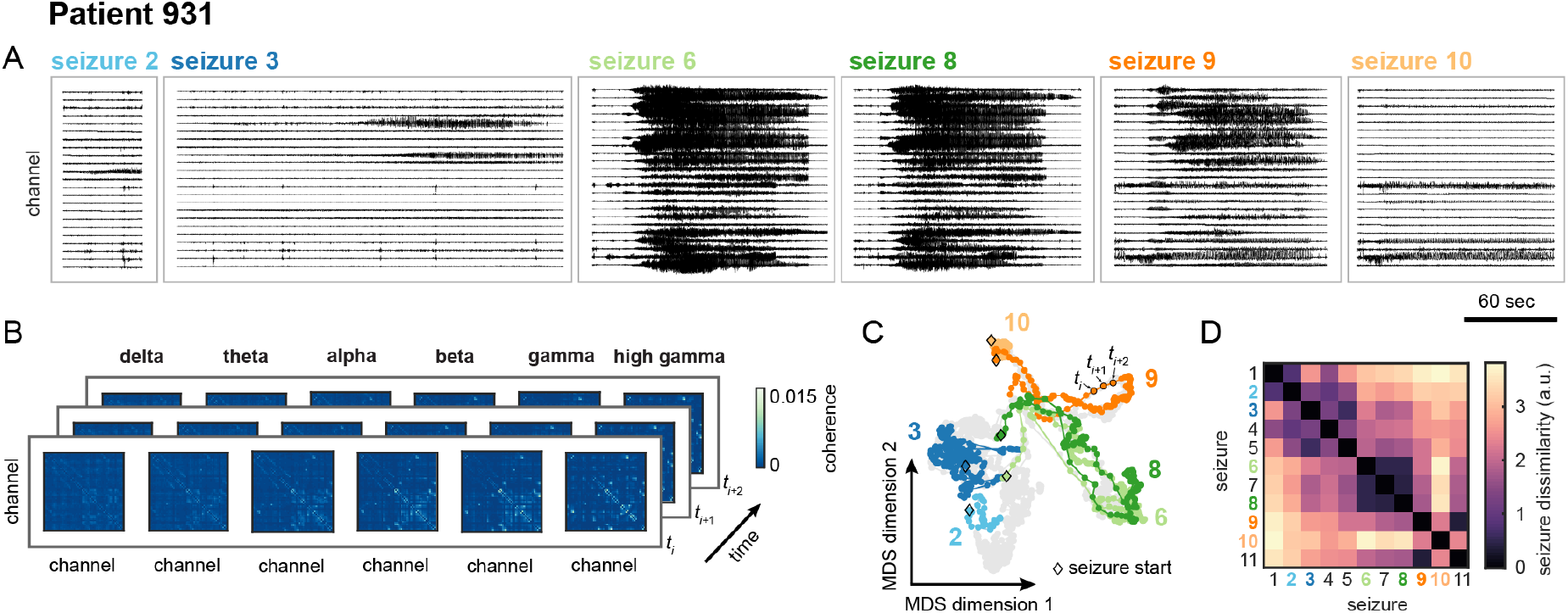
Visualising and comparing seizure pathways through network space in an example patient, patient 931. A) Intracranial EEG traces of a subset of the patient’s seizures. For clarity, only a representative subset of the recording channels are shown. B) Functional connectivity of three example seizure time windows. Functional connectivity was defined as band-averaged coherence in each of six different frequency bands. Each matrix was normalised so that the upper triangular elements summed to one. Self connections are not shown in order to focus on inter-channel connectivity. C) Projection of all seizure time windows into a two dimensional space using multidimensional scaling (MDS), allowing visualisation of seizure pathways through network space. Each point corresponds to a seizure time window, and time windows with more similar network dynamics are placed closer together in the projection. Consecutive time windows in the same seizure are connected to visualise seizure pathways. The time windows and pathways of the six seizures shown in Fig. 1A have been highlighted using the corresponding colours, and the time windows of the remaining seizures are shown in grey for reference. The first time windows of the selected seizures are each marked with a diamond. D) Seizure dissimilarity matrix of all of the patient’s seizures, which quantifies the difference in the network dynamics of each pair of seizures. A low dissimilarity indicates that the two seizures have similar pathways through network space.

Due to the high dimensionality of this network space, it was infeasible to directly visualise seizure pathways. However, seizure pathways could be approximated in a two dimensional projection using multidimensional scaling (MDS), a dimensionality reduction technique that attempts to maintain the distances between high-dimensional data points in the lower dimensional space (Fig. 1C). As such, this technique placed seizure time windows in the two-dimensional projection based on the similarity of their network configurations; each time window was represented by a single point, and points corresponding to time windows with more similar network dynamics were placed closer together. While imperfect, this approximation of the network space nonetheless provided an intuitive visualisation for comparing seizure pathways in the same patient. For example, in patient 931, the projection demonstrated that two seizures may follow approximately the same pathway (seizures 6 and 8), part of the same pathway (seizures 8 and 9), or completely distinct pathways (seizures 2 and 10) through the network space, in agreement with visual impressions of the EEG.

To quantify these visual observations, we developed a “seizure dissimilarity” measure that provides a “distance” between two seizures based on their pathways through network space. Importantly, our approach recognises similarities in seizure pathways, even if the seizures evolve at different rates, by first applying dynamic time warping (44) to each pair of seizure functional connectivity time courses (SI Appendix, Text S2). Dynamic time warping nonlinearly stretches each time series such that similar points are aligned, thus minimizing the total distance between the two time series. We then defined the dissimilarity between two seizures as the average difference between the seizure pathways across all warped time points. The seizure dissimilarity matrix then summarises the dissimilarity between all pairs of seizure pathways in the same patient (Fig. 1D). In patient 931, seizures with similar pathways therefore have a low dissimilarity (e.g., seizures 6 and 8, dissimilarity 0.49); seizures with distinct, distant pathways have high dissimilarity (e.g., seizures 2 and 10, dissimilarity 3.21); and seizures with partially overlapping pathways have an intermediate level of dissimilarity (e.g., seizures 8 and 9, dissimilarity 1.75). Again, our measure of seizure dissimilarity agrees with intuitive comparisons of seizures based on visually assessing the iEEG (Fig. 1A) and MDS projections of the seizure pathways (Fig. 1C).

It is important to note that both seizure dissimilarity matrices and MDS projections are patient-specific: due to different electrode implantations, we cannot compare seizures across patients using these network features. However, because we normalise the magnitude of the functional connectivity, we can compare seizure dissimilarity values across patients, even if the patients have different numbers of recording electrodes. In the remainder of the paper, we will focus on the across patient results, while using patient 931’s seizures as examples. The seizure variability analysis of all patients will be available on Zenodo (http://dx.doi.org/10.5281/zenodo.3560736) and summarised in SI Appendix, Text S3.

### Seizure variability is a common feature in all patients

Using our measure of seizure dissimilarity, we compared seizure pathways through network space in each patient. We first determined if seizure variability was present in all patients. Fig. 2A shows the distribution of seizure dissimilarities in each patient, with patients sorted from lowest (patient 934) to highest (patient I002 P006 D01) median dissimilarity. Note that each point corresponds to the difference in network dynamics of a *pair* of seizures, rather than a feature of a single seizure. From these distributions, it is apparent that all patients had variability in seizure network dynamics. Even in patients with more consistent seizures, such as patient 934, there were pairs of seizures with high dissimilarity, indicating dissimilar seizure pathways. Meanwhile, other patients, including patient 931, had varying levels of different dynamics, with only a few pairs of similar seizures.

**Fig. 2:**
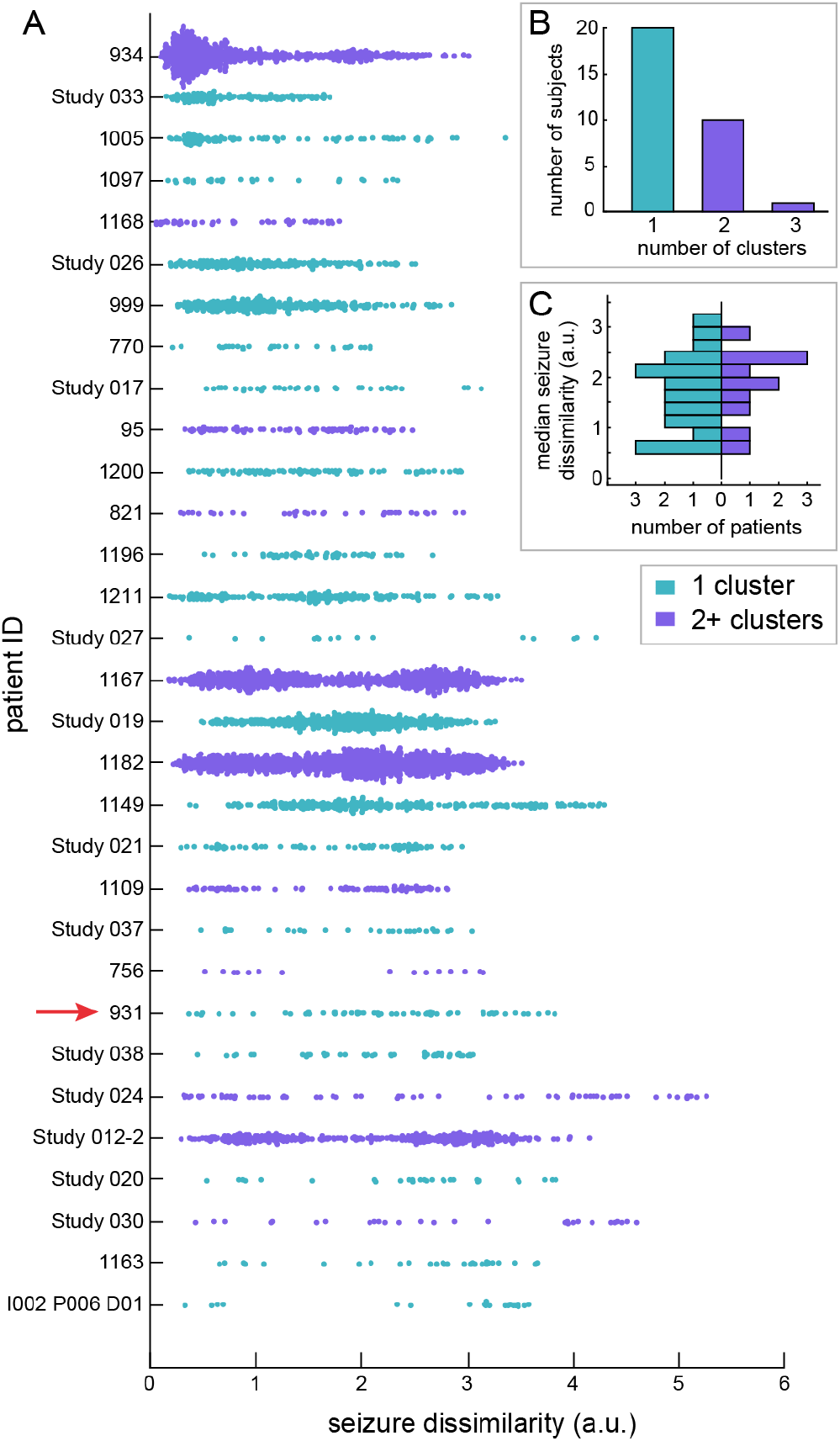
Variability in seizure pathways is common in all patients, but may take the form of either a spectrum or clusters of seizure dynamics. A) Distributions of seizure dissimilarities in each patient. Patients are sorted from lowest median seizure dissimilarity (patient 934) to highest median seizure dissimilarity (I002 P006 D01). The red arrow indicates patient 931, the example patient from Figure 1. In each distribution, each point corresponds to the dissimilarity of a pair of seizures. The distribution is coloured based on the number of seizure clusters, computed using seizure dissimilarities, in each patient. B) The number of patients with different numbers of seizure clusters based on seizure dissimilarities. The majority of patients had one seizure cluster. C) Distribution of median seizure dissimilarities in patients with one (left, teal) or multiple (right, purple) seizure cluster.

Past studies have noted that some patients have populations of seizures with distinct features such as different onset sites (9, 11) or durations (8, 12). As such, we would expect the variability described in these studies to result from different, discrete seizure types coexisting in the same patient. We therefore tested if each of the patients in our cohort had multiple seizure types by clustering their seizures based on seizure dissimilarities (Fig. 2B; see Methods for details). Contrary to our expectation, we found that the majority of patients (21 patients), including patient 931, did not have distinct types. Importantly, without a clear way to split their seizures into different types, the full diversity of their seizure dynamics could not be described by a few example seizures. Ten patients had two or more seizure clusters, although there was still variability in dynamics within most clusters (SI Appendix, Text S4), and the average amount of seizure variability was the same in patients with or without multiple seizure clusters (Fig. 2C) (two sample *t*-test, *p* = 0.68). Thus, the presence or absence of different types of seizure dynamics does not indicate the average amount of seizure variability in each patient.

We also found that the observed variability was not solely explained by the presence of different clinical seizure types (subclinical, focal, or secondarily generalised seizures) (SI Appendix, Text S5). This finding was expected given that seizures of different clinical types can share similar dynamics, while seizures of the same clinical type may have dramatically different features (16, 45, 46). Additionally, we found no association between postsurgical seizure freedom and measures of seizure variability (SI Appendix, Text S6). Likewise, higher levels of seizure variability were not associated with a particular seizure onset site (SI Appendix, Text S6). These findings suggest that the level of seizure variability is not associated with certain patient pathologies or treatment outcomes; instead, other factors may be more crucial for determining the extent and form of the variability.

### Seizures with more similar pathways tend to occur closer together in time

Many time-varying factors, such as sleep (21, 23, 45, 47, 48) and hormones (49–52), are thought to influence seizure likelihood and dynamics. Additionally, during presurgical monitoring, antiepileptic medication is reduced in many patients, impacting brain dynamics (53). We therefore explored whether there is a temporal structure to how seizure dynamics change over time in each patient. Fig. 3A shows the pathways of patient 931’s seizures, as well as the time that each seizure occurred relative to the patient’s first seizure. From this visualisation, we see that the pathways gradually migrated through network space as the recording progressed, creating the observed spectrum of network dynamics. Moreover, looking at the seizure timings, we also see that seizures with similar pathways, such as seizures 6-8, tended to occur close together in time.

**Fig. 3:**
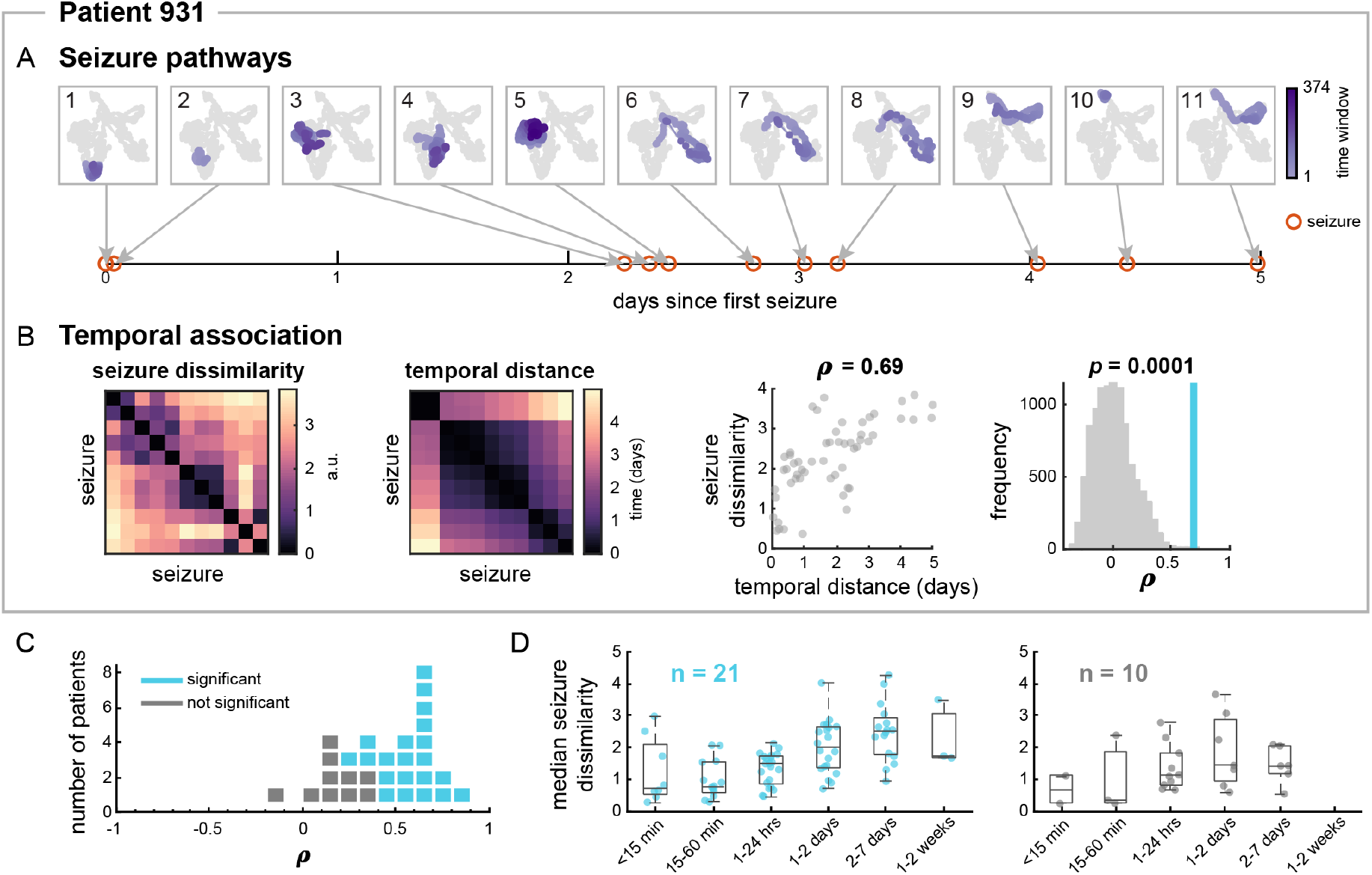
More similar seizures tend to occur closer together in time in most patients. A) MDS projections of all of patient 931’s seizure pathways, numbered from first to last seizure. The pathway of each seizure is shown in purple, with earlier time windows in lighter purple. The time windows and pathways of the remaining seizures are shown in grey for comparison. Below the pathways, the time of each seizure (red circles) relative to the first seizure is shown. Note that seizures with more similar pathways tend to occur close together in time. B) From left to right: patient 931’s seizure dissimilarity matrix, temporal distance matrix, and comparison of seizure dissimilarities and temporal distances. The temporal distance matrix quantifies the amount of time between each pair of seizures, in days. Plotting the seizure dissimilarity vs. the corresponding temporal distance of each pair of seizures (scatter plot, second from right) reveals a positive Spearman’s correlation *ρ* between the two features. The significance of this correlation can be tested using permutation testing (distribution, far right). The distribution of the 10,000 correlations computed from permuted matrices is shown in grey, and the observed correlation is marked with the vertical blue line. The *p*-value of the association was equal to the proportion of times a correlation value greater than or equal to the observed correlation was seen in the distribution. C) Dot plot showing the range of correlations between seizure dissimilarities and temporal distances across all subjects. Each marker represents a patient (blue = significant correlation, grey = not significant after false discovery rate correction). D) Median seizure dissimilarities of pairs of seizures occurring within different time intervals (i.e., temporal distances) for patient with (left, blue) and without (right, grey) a significant correlation between seizure dissimilarities and temporal distances. Each point corresponds to the median dissimilarity of pairs of seizures occurring within the given time interval in a single patient. Note that some time intervals have fewer observations since some temporal distances were not observed in some subjects. The boxplots indicate the minimum, lower quartile, median, upper quartile, and maximum of the distribution of median seizure dissimilarities, across the subset of subjects, for that time interval.

To quantify this temporal relationship, we defined a “temporal distance” matrix as the amount of time elapsed between each pair of the patient’s seizures (Fig. 3B). Patient 931’s seizure dissimilarity and temporal distance matrices have strikingly similar structures: groups of seizures with low dissimilarity tended to occur together in a relatively short time interval. In this patient, there was a strong and significant positive correlation between these features (Spearman’s ρ = 0.69, *p* = 0.001, one-tailed Mantel test), indicating that seizures with more similar pathways tended to occur closer together in time.

Fig. 3C summarises the relationship between seizure dissimilarities and temporal distances across all patients. In almost all patients, there was a positive Spearman’s correlation between seizure dissimilarities and temporal distances (range: −0.10 – 0.83, mean: 0.45). This association was significant in 21 patients (67.7%) after false discovery rate correction. In these patients, we also observed that the average level of dissimilarity tends to increase with the time between the two seizures (Fig. 3D). Interestingly, there was no association between whether antiepileptic medication was reduced and whether the correlation between seizure dissimilarities and temporal distances was significant (χ^2^ test, *p* = 0.96) (SI Appendix, Text S7). Therefore, although medication levels may affect seizure dynamics (9, 16, 54, 55), medication changes alone cannot explain the observed shifts in seizure pathways, suggesting that other temporal factors also play a role in shaping seizure features.

### Seizure pathways change on different timescales

The observed temporal associations of seizure dissimilarities reflected gradual changes in seizure dynamics across the length of each recording. In other words, we observed relatively slow shifts in seizure pathways over the course of multiple days. However, we also hypothesised that seizure dynamics may change on shorter timescales due to, for example, circadian rhythms. Such rhythms would create timescale-dependent relationships between seizures; in particular, there would be a positive correlation between seizure dissimilarities and temporal distances on shorter timescales, but this association would be destroyed on longer timescales.

Therefore, to explore the possibility of different timescales of changes in seizure dynamics, we scanned the correlation between seizure dissimilarities and temporal distances on different timescales *T* ranging from 6 hrs to the longest amount of time between a seizure pair (Fig. 4A). For example, for *T* = 3 days, we computed the correlation between seizure dissimilarities and temporal distances for all pairs of seizures that occurred within three days of each other. We refer to these sets of correlation as “temporal correlation patterns” of seizure dynamics. Fig. 4A shows the temporal correlation patterns of patient 931’s seizures. As we determined earlier, there was a positive correlation between seizure dissimilarities and temporal distances when all seizures were included in the computation (*T* = 5 days) as a result of the observed gradual changes in seizure pathways. At shorter timescales, however, the temporal relationship fluctuates; for example, the correlation is relatively low at *T* = 1 and 2.5 days, and higher at *T* = 0.75 and 2.5 days. These fluctuations are signs of additional, timescale-dependent changes in seizure dynamics beyond the gradual changes.

**Fig. 4:**
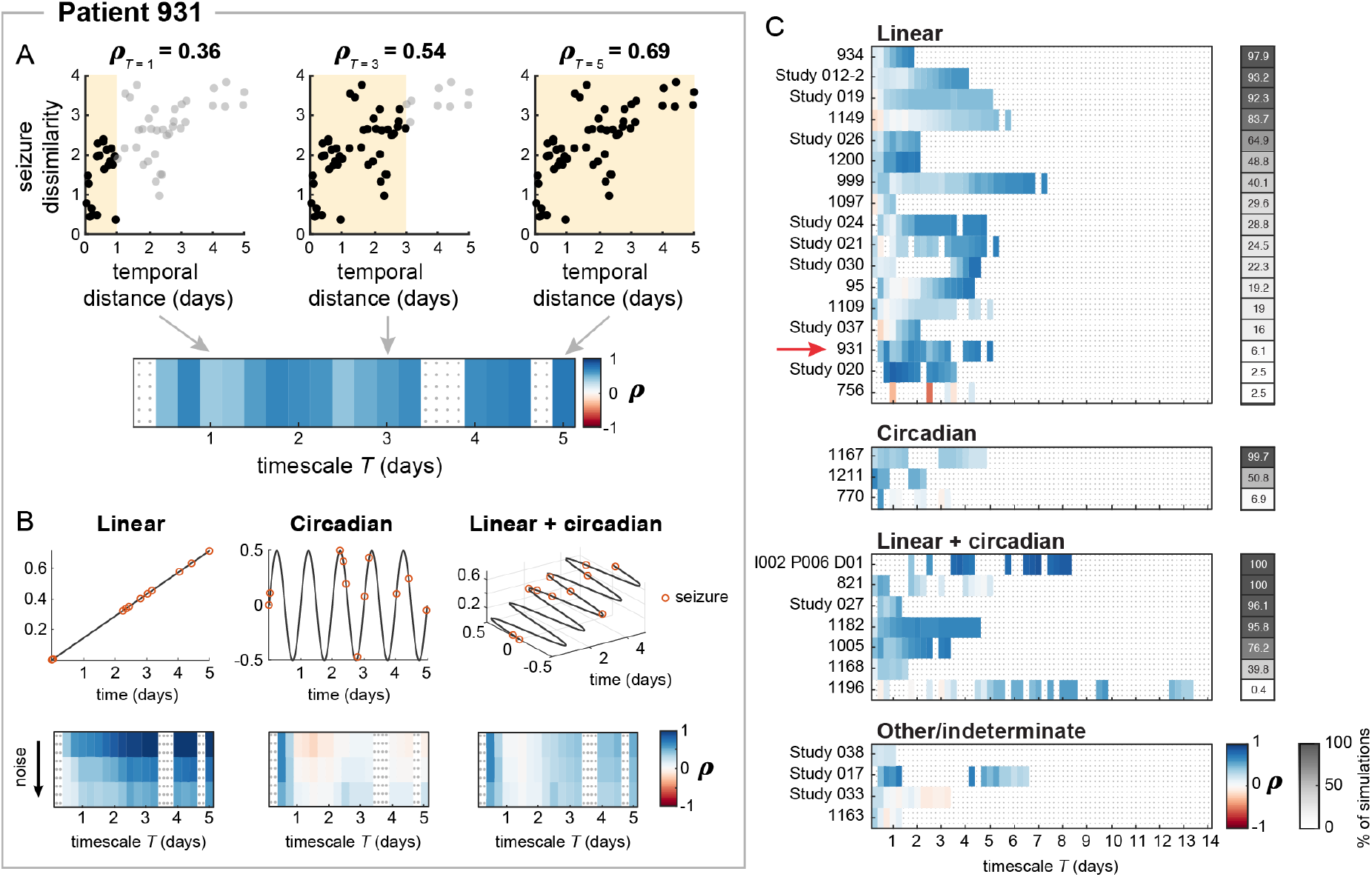
Temporal patterns of changes in seizure dynamics. A) For patient 931, the correlation between seizure dissimilarities and temporal distances was computed for seizure pairs within different timescales, producing a heatmap of the “temporal patterns” of seizure dynamics (bottom). The seizure pairs used to compute the correlation for three example timescales (*T* = 1 day, *T* = 3 days, and *T* = 5 days) are shown in the top scatter plots (reproduced from Fig. 3B). Purple shading indicates the timescale used for each computation (e.g., seizure pairs occurring within 0 – 1 days for *T* = 1 day), black points correspond seizure pairs used to compute the correlation for that timescale, and grey points correspond to seizure pairs occurring further apart than the given timescale. The correlation between seizure dissimilarities and temporal distances at the given timescale is shown above each scatter plot. At *T* = 5 days, all seizure pairs are included in the computation, producing the same temporal correlation as in Fig. 3B. If there were less than seven seizure pairs occurring within a given timescale, or if no new seizure pairs were added when the timescale was extended, the correlation for that timescale was excluded from the heatmap and downstream analysis (regions with grey dots). B) Seizure dissimilarities were modelled based on linear (left), circadian (middle) or a combination of linear + circadian (right) changes in seizure dynamics. The timepoints of patient 931’s seizures are shown in red on each function. From each model, the temporal pattern of seizure changes was then derived (heatmaps, bottom row), revealing the expected temporal associations between seizures on different timescales given the simulated changes in dynamics. The temporal pattern also depended on the amount of noise included in the simulation; for clarity and brevity, different levels of a single noise realisation are shown, with the amount of noise increasing from the top to bottom row of each set of heatmaps. C) Temporal patterns of seizure dynamics in each patient, sorted by the type of model that most closely matched the observed temporal patterns. The heatmap on the right (grey) shows the percentage of noisy simulations of the selected parameter set that closely matched the observed dynamics. Patient 931 is indicated with a red arrow.

To investigate how these temporal correlation patterns arise, we modelled different patterns of seizure variability and the corresponding temporal correlation patterns (Fig. 4B) (see Methods and SI Appendix, Text S8 for modelling details). Specifically, for each patient, we simulated sets of seizure dissimilarities arising from different levels of linear, circadian, and/or noisy dynamics based on predefined time-varying functions and the patient’s seizure times. Linear changes in dynamics would correspond to the slower, gradual shifts in seizure dynamics; circadian changes represent dynamics modulated by circadian rhythms; and noisy changes allow for the influence of random fluctuations and intermittent factors. From these simulated dissimilarities, we computed simulated temporal correlation patterns. Based on the model parameters that most reliably reproduced the observed temporal correlation pattern, we categorised each patient’s pattern of seizure variability as linear (Fig. 4B, left), circadian (Fig. 4B, middle), or linear + circadian (Fig. 4B, right). Crucially, this modelling approach allowed us to hypothesise how different patterns of seizure variability could interact with the patient’s seizure timings to produce the observed temporal relationships between seizures.

In patient 931, example simulations using a single noise realisation demonstrated that these different underlying models could produce different temporal correlation patterns of seizure dynamics (Fig. 4B). A linear change in seizure dynamics produced a positive temporal relationship that is stronger at longer timescales. Higher levels of noise reduced this positive correlation at all timescales. Meanwhile, a circadian model only produced strong, positive temporal correlations at timescales shorter than one day. Finally, a combination of the linear and circadian factors created both the short-term temporal relationships and a positive temporal correlation at the longer timescales. Note that there were also some additional fluctuations in the temporal correlation patterns due to noisy changes in dynamics, especially at higher levels of noise, which will differ depending on the outcome of the noisy simulation.

Fig. 4C shows the underlying model (linear, circadian, or linear + circadian) most likely to underlie the observed temporal correlation patterns, as defined by the percentage of model simulations with matching temporal correlation patterns. We additionally required the selected model to 1) outperform noisy simulations alone, 2) clearly distinguish between the linear and circadian models, and 3) in the case of the linear + circadian model, clearly outperform one of the simpler models. Using these criteria, seventeen patients’ temporal correlation patterns were best explained by the linear model, three by the circadian model, and seven by the linear + circadian model. Thus, most patients (77.4%) required a linear component to explain the observed changes in seizure dynamics, while (32.3%) of the temporal correlation patterns were well-matched by a model incorporating circadian dynamics. As before, different classifications of seizure dynamics were not associated with surgical outcomes (SI Appendix, Text S6) or whether the patient’s medication was reduced during presurgical monitoring (SI Appendix, Text S7).

Four patients’ temporal correlation patterns could not be assigned to a model, either because the linear model and circadian model performed similarly (patient Study 038) or the best model did not outperform noise alone (patients Study 017, Study 033, and 1163). Notably, in some patients (Study 020, 756, 1196, and Study 017) only a small percentage of the simulations matched that observed temporal correlation patterns, indicating that reproducing the observed dynamics required specific patterns of noise. In these cases, other models may therefore provide a better explanation for the patient’s changes in seizure dynamics. In particular, many of these patients had strong positive correlations at a timescales longer than one day, but less than the length of the recording, suggesting multi-day fluctuations in seizure dynamics.

## Discussion

We have quantified variability in seizure network dynamics within individual human patients with focal epilepsy, revealing that within-patient seizures are neither deterministic nor comprehensively represented by a single dynamical pathway. Contrary to our expectation, most patients had a spectrum of seizure dynamics, rather than distinct seizure populations. Interestingly, seizure network dynamics change over time in most patients, with more similar seizures tending to occur closer together in time. Our modelling results indicate that in most patients, a combination of fast (i.e. circadian) and/or slow changes in seizure pathways may underlie the observed variability, suggesting that factors operating on different timescales modulate within-patient seizure dynamics.

We investigated variability in seizure functional network evolution due to the importance of network interactions in ictal processes (2, 7, 22, 31, 33–43) and build on previous work by demonstrating within-patient variability in these pathological network dynamics. However, the framework we present could easily be adapted to compare other features that highlight different aspects of seizure dynamics. For example, a univariate feature that captures the amplitude and frequency of ictal discharges may be better suited for comparing the involvement of different channels, similar to how clinicians visually compare EEG traces. Meanwhile, comparisons of parameter time courses, derived using model inversion (8, 56, 57), could reveal different patterns of changes in the neural parameters underlying a patient’s seizures. Finally, due to patient-specific recording layouts, we focused on comparing seizure dynamics within individual patients. However, comparing seizures across patients, either using spatially-independent features or common recording layouts, in future studies could uncover common classes of pathological dynamics (8, 58).

To quantify within-patient variability in seizure pathways, we developed a “seizure dissimilarity” measure that addresses the challenges of comparing diverse spatiotemporal patterns across seizures. A few previous studies have attempted to quantitatively compare seizure dynamics using either univariate (26, 27, 29, 30) or network (25, 28) features computed from scalp or intracranial EEG. These earlier dissimilarity measures were based on edit distance, which captures how many replacements, insertions, and deletions are required to transform one sequence into another. Importantly, the insertion cost increases the dissimilarity of similar seizures with different rates of progression. Although previous work suggested lowering seizure dissimilarity in such scenarios (30), to our knowledge, our dynamic time warping approach provides the first measure of seizure dissimilarity that does not penalise temporal variability between otherwise similar seizures. Despite this difference, those past studies also reported both common and disparate dynamics across within-patient seizures; however, their analysis was limited to a small number of patients and/or seizures per patient. Our work provides novel insight into the prevalence and characteristics of seizure variability by analysing over 500 seizures across thirty-one patients. Finally, we expand on previous work by using seizure dissimilarity for downstream analysis, including clustering seizures and describing temporal changes in seizure dynamics.

Previous work has found that within-patient seizures have similar dynamics (2–8), although variability may be introduced through different rates of progression (4, 59) or early termination in the seizure pathway (6, 8). In our cohort, we observed that subsets of within-patient seizures follow approximately the same dynamical pathway through network space, and such similar groups of seizures likely underlie these past findings. However, we also found that the complete repertoire of within-patient seizure network dynamics is poorly characterised by a single, characteristic pathway. Notably, we also found that a patient with different seizure dynamics does not necessarily have distinct populations of seizures. We therefore propose a model in which various decision points, existing on the framework of potential seizure pathways, produce a repertoire of seizure evolutions (SI Appendix, Text S9). The number and location of these decision points would also explain why some patients have a spectrum of seizure dynamics: a larger number of “forks” in seizure pathways would produce a series of small changes between different seizures, rather than distinct seizure types. Future studies can map these potential seizure pathways and the factors shaping how individual seizures evolve.

The crucial question is then how these different seizure pathways arise from the same neural substrate. In theory, a range of changes before or during the seizure can affect its network progression. We hypothesise that spatiotemporal changes in the interictal neural state produce seizures with different characteristics. Past studies suggest that neural excitability (19, 55, 60), inhibition (59), and network interactions (22, 61) influence certain spatiotemporal seizure features, such as the rate and extent of seizure propagation. These changes in brain state may be driven by various factors, including sleep (21, 45, 47), hormones (49–52), and medication (53). Recently, prolonged recordings of patients with focal epilepsy have revealed that the rates of epileptiform discharges and seizures fluctuate according to both circadian and patient-specific multidien (approximately weekly to monthly) cycles (48, 62). An intriguing possibility is that the same factors that rhythmically modulate seizure likelihood may also influence seizure dynamics. Consistent with this hypothesis, we found that the majority of observed temporal patterns of seizure variability were well-explained by models incorporating circadian and/or linear changes in seizure dynamics. In particular, the linear component of the model may reflect gradual changes in dynamics on slower timescales, ranging from weeks to months. These simple models provided an initial hypothesis for the observed patterns of changes in seizure dynamics. Some patients seizure patterns may be better explained by more complex models that capture different dynamics, such as multistability or multidien cycles. Ultimately, it is likely that various factors, with differential effects on seizure dynamics, interact to produce the observed repertoire of seizure network evolutions. Analysing within-patient seizure variability in long-term recordings could provide additional insight into patterns of temporal changes in seizure dynamics.

Notably, a large number of the patients in our study underwent antiepileptic medication reduction as part of pre-surgical monitoring, making it difficult to disentangle the effects of changing drug levels from other potential slow-varying modulators of seizure dynamics. Changes in antiepileptic medication can impact neural excitability (63–65), and medication tapering increases seizure likelihood in most patients (16, 66); however, it is controversial whether it also affects seizure patterns (9, 16, 54, 66). In some cases, it appears that medication tapering reveals latent seizure pathways that are suppressed by medication (9) or allows existing pathways to further progress (e.g., the secondary generalisation of typically focal seizures) (16). It is possible that the impact of medication reduction on seizure dynamics is drug-, patient-, and dose-dependent, and may ultimately depend on how well the medication controls neuronal excitability (55). However, medication changes alone cannot account for the observed seizure variability in our cohort, as we observed temporal associations of seizure dynamics in patients that did not undergo medication reduction. In future work, associating medication levels with differences in seizure dynamics could help untangle the different factors shaping seizure dynamics.

Another confounding factor in our data is that the surgical implantation itself could artificially alter seizure dynamics. Using chronic recordings of epileptic canines, Ung *et al.* (67) found variability in seizure onset and interictal burst dynamics, with the most stable dynamics emerging approximately a few weeks after electrode implantation. In agreement with their work, we found that earlier seizure types often recur later in the recording, making it unlikely that gradual changes in the recording quality or an acute reaction to the surgery underlie the observed variability. Instead, Ung *et al.* hypothesised that seizure variability results from transient, atypical dynamics as the brain recovers from surgery, with later dynamics representing a truer epileptic network. Other stressors, such as medication withdrawal, could similarly elicit abnormal dynamics. Nevertheless, a large number of our patients had good surgical outcomes, suggesting that their recorded seizures accurately represented their epileptic networks. Additionally, clinicians often note that patients have typical seizures during iEEG recordings, as compared to preimplantation reports, despite the effects of surgery and medication withdrawal (16). As such, the observed seizure dynamics in our cohort may be part of their usual repertoires of seizure dynamics, even if some dynamics are only elicited by strong stressors. Further analysis in chronic human recordings is needed to determine whether and how seizure pathways vary in a more naturalistic setting.

Contrary to the expectation that high levels of seizure variability may worsen surgical outcomes, we found no association between these patient features. It may be that only some types of variability, such as multifocal (9) or secondarily generalised (68) seizures, impact the likelihood of seizure freedom following surgery. Importantly, variability in the seizure onset network state does not indicate that a patient has multifocal seizures, as different network configurations can be associated with the same apparent ictal onset zone. Additionally, variability in seizure dynamics may not be inherently deleterious, as long as it is observed and accounted for when planning the surgical resection. Indeed, due to the short presurgical monitoring time and limited spatial coverage of the recording electrodes, some potential seizure pathways may not have been captured (11, 67), leading us to underestimate the level of variability in some patients.

Although the amount of seizure variability was not associated with post-surgical seizure freedom, it may have implications for clinical treatments. First, regardless of the source of the observed seizure variability, the different seizure dynamics observed during presurgical monitoring provide crucial information for guiding surgical resection. For example, recent studies suggest that seizure network properties can help identify epileptogenic tissue (7, 69, 70); however, we must determine if seizures with different network evolutions provide equivalent localisation information. Seizure variability may also have implications for seizure prediction. In particular, in that same patient, seizures with different dynamics may have distinct preictal signatures, making seizure prediction more difficult (10, 12). A successful seizure prediction algorithm would either need to recognise multiple signatures or find common features among the disparate preictal dynamics. Finally, neurostimulation offers a promising new approach for controlling seizures; however, in rodent models, the effectiveness of a given stimulation protocol depends on the preictal brain state (18). Thus, such interventions may need to recognise and adapt to the specific characteristics of each seizure type in order to control all seizure dynamics. Importantly, our cohort was limited to patients with medication refractory focal epilepsy who were candidates for surgical resection. The characteristics and clinical implications of seizure variability may be different in other patient cohorts.

In summary, we have shown that there is within-patient variation in seizure network dynamics in patients with focal epilepsy. Temporal changes in seizure dynamics suggest that a combination of circadian and slow-varying factors shape these seizure pathways, perhaps by modulating the background brain state. Further research is needed to determine whether and how preictal dynamics shape seizure pathways. Uncovering these mechanisms could provide novel approaches for predicting and controlling seizures that are tailored to the complete repertoire of pathological neural dynamics in each patient.

## Materials and methods

### Patient selection and data acquisition

This work was a retrospective study that analysed seizures from 13 patients from the Mayo Clinic and the Hospital of the University of Pennsylvania (available on the IEEG Portal, www.ieeg.org (71, 72)) and 18 patients from the University College London Hospital (UCLH) who were diagnosed with refractory focal epilepsy and underwent presurgical monitoring. Patients were selected without reference to cause or other characteristics of their pathology. All IEEG Portal patients gave consent to have their anonymised iEEG data publicly available on the International Epilepsy Electrophysiology Portal (www.ieeg.org) (71, 72). For the UCLH patients, their iEEG was anonymised and exported, and the anonymised data was subsequently analysed in this study under the approval of the Newcastle University Ethics Committee (reference number 6887/2018).

For each patient, the placement of the intracranial electrodes was determined by the clinical team, independent of this study. Ictal segments were identified and extracted for the analysis based on clinical seizure markings. To be included in the study, each patient was required to have had at least six seizures suitable for the analysis. This threshold was chosen to allow examination of seizure variability in a broad cohort of subjects, while still ensuring that enough seizures were observed to draw conclusions about the forms, types, and characteristics of seizure variability in each subject. Seizures were excluded from the analysis if they did not have clear electrographic correlates (with clear onset and termination), if they were triggered by/occurred during cortical stimulation, if they had noisy segments, or if they had large missing segments. Periods of status epilepticus and continuous epileptiform discharges were also excluded. However, electrographic seizures without clinical correlates were included in the analysis. Additional information about each subject and the analysed seizures is shown in SI Appendix, Text S1.

### iEEG preprocessing

For each patient, if different seizures were recorded at multiple sampling frequencies, all of the recordings were first downsampled to the lowest sampling frequency. Noisy channels were then removed based on visual inspection. In the remaining channels, short sections of missing values were linearly interpolated. These sections of missing values were <0.05 s with the exception of one segment in seizure 2 of patient “Study 020”, which was 0.514 s. All channels were re-referenced to a common average reference. Each channel’s time series was then bandpass filtered from 1-150 Hz (4^th^ order, zero-phase Butterworth filter). To remove line noise, the time series were additionally notch filtered (4^th^ order, 2 Hz width, zero-phase Butterworth filter) at 60 and 120 Hz (IEEG Portal patients) or 50, 100, and 150 Hz (UCLH patients).

### Computing functional connectivity

To compute the time-varying functional connectivity of each seizure, a 10 s sliding window, with 9 s overlap between consecutive windows, was applied to each preprocessed ictal time series. The same sliding window parameters have previously been used to estimate time-varying coherence in ictal iEEG data (73). For each window, the coherence between each pair of iEEG channels was computed in six different frequency bands (delta 1-4 Hz, theta 4-8 Hz, alpha 8-13 Hz, beta 13-30 Hz, gamma 30-80 Hz, high gamma 80-150 Hz). The coherence in each frequency band was computed using band-averaged coherence, defined as

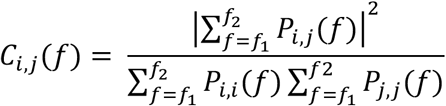

where *f*_1_ and *f*_2_ are the lower and upper bounds of the frequency band, *P*_*i,j*_(*f*) is the cross-spectrum density of channels *i* and *j*, and *P*_*i,i*_(*f*) and *P*_*j,j*_(*f*) are the autospectrum densities of channels *i* and *j*, respectively. In each window, channel auto-spectrums and cross-spectrums were calculated using Welch’s method (2 s sliding window with 1 s overlap).

Thus, in a patient with *n* iEEG channels, the functional connectivity of each time window was described by six symmetric, non-negative, *n*×*n* matrices, in which each entry (*i*,*j*) gives the coherence between channels *i* and *j* in the given frequency band. Each matrix was then written in vector form by re-arranging the upper-triangular, off-diagonal elements into a single column vector of length (*n*^2^ – *n*)/2. Each vector was then normalised so that the *L*1 norm (i.e., sum of all elements) was 1, thus ensuring that differences between connectivity vectors captured a change in connectivity pattern rather than gross changes in global levels of coherence. This normalisation step also allowed the magnitude of seizure dissimilarities to be compared across patients with different numbers of electrodes. For each time window, the six connectivity vectors were then vertically concatenated together, forming a single column vector of length 6*(*n*^2^ – *n*)/2. Each patient’s ictal connectivity vectors were subsequently horizontally concatenated together to form a matrix *V* containing 6*(*n*^2^ – *n*)/2 features and *m* observations, where *m* is the total number of ictal windows across all seizures.

### Dimensionality reduction and visualisation

Small fluctuations in the functional connectivity due to noise would create a high baseline dissimilarity between seizures. Therefore, to reduce noise in the connectivity matrices, non-negative matrix factorization (NMF) (74) was used to approximately factor each patient’s ictal time-varying connectivity matrix *V* into two non-negative matrices, *W* and *H*, such that *V*≈*W*×*H* (details provided in SI Appendix, Text S10). The matrix *W* contained patient-specific basis vectors, each of which had 6*(*n*^2^ – *n*)/2 features that captured a pattern of connectivity across all channels and frequency bands. Each original ictal time window was summarised as an additive combination of these basis vectors, with the coefficients matrix *H* giving the contribution of each basis vector to each time window. These factorisations were patient-specific since the basis vector features depended on the iEEG electrode layout in each patient. The optimal number of basis vectors, *r*, was determined using stability NMF (75).

For each patient the selected factorisation was then used to create *V*=W*×*H*, a lower-rank approximation of the original time-varying seizure functional connectivity (SI Appendix, Text S10). This return to the original feature space is necessary since NMF basis vectors are not orthogonal, and distances in NMF basis vector space are therefore not equivalent to distances in feature space. Each reconstructed connectivity vector was then re-normalised to have an *L*1 norm of 1, ensuring that differences in reconstruction accuracy did not affect the distances between different ictal timepoints. To visualise the connectivity vectors of patient 931’s seizures in Fig. 1C, all time seizures windows were projected into a two-dimensional embedding using multidimensional scaling (specifically, Sammon mapping) based on their *L*1 (cityblock) distances in the high-dimensional reconstructed feature space.

### Computing seizure dissimilarities

Following the NMF-based reconstruction of the seizure connectivity, the network evolution of each seizure was described by a multivariate time series with 6*(*n*^2^ – *n*)/2 features. To compare network evolutions across within-patient seizures, a “seizure dissimilarity matrix” was created for each patient. Each pair of seizure functional connectivity time series was first warped using dynamic time warping, which stretches each time series such that the total distance between the two time series is minimised (SI Appendix, Text S2). This step ensures that 1) similar network dynamics of the two seizures are aligned, and 2) the warped seizures are the same length. We chose to minimise the *L*1 distance between each pair of seizures, as this metric provides a better measure of distances in high-dimensional spaces (76).

Following dynamic time warping, the *L*1 distance between the pair of warped time series was computed, resulting in a vector of distances capturing the dissimilarity in the seizures’ network structures at each time point. The “seizure dissimilarity” between the two seizures was defined as the average distance across all warped time points. The seizure dissimilarity matrix contains the dissimilarities between all pairs of the patient’s seizures. Note that seizure dissimilarity is not a metric distance because the triangle equality does not necessarily hold; however, it performs similarly to alternative metric distances of seizure dissimilarity (SI Appendix, Text S11).

### Seizure clustering and cluster evaluation

To identify groups of similar seizures in each patient, each patient’s seizures were hierarchically clustered by using the seizure dissimilarity matrix as input for an agglomerative hierarchical clustering algorithm, UPGMA (unweighted pair group method with arithmetic mean). The hierarchical clustering resulted in a dendrogram that summarised the similarity between the patient’s seizures. Note that the hierarchical clustering representation was an approximation of the seizure dissimilarities that forced all dissimilarities into a metric space.

The gap statistic (77), which compares the within-cluster dispersion of a given clustering relative to a reference (null) distribution, was then used to determine if optimal flat (i.e., non-hierarchical) clusters of seizures existed in each patient. In order to generate reference datasets, the patient’s seizures were first projected into Euclidean space using classical (Torgerson’s) multidimensional scaling (MDS). Note that this step differs from the earlier visualisation of seizure pathways, which projected seizure time points, rather than seizures themselves. Given the seizure dissimilarity matrix, MDS assigned a coordinate point to each seizure while attempting to preserve the specified dissimilarities between seizures. In order to most closely approximate the dissimilarities matrix, the seizures were projected onto the maximum possible number of dimensions; note, however, that like the hierarchical clustering, MDS also provided a metric approximation of the nonmetric dissimilarities. One thousand reference datasets were then generated by drawing coordinates from a uniform distribution placed over a box aligned with the principal components of the projected seizure data. Each reference dataset was hierarchically clustered by computing the distances between the coordinate points and applying the UPGMA algorithm. To test for flat clusters in the seizure data and reference datasets, the dendrograms were cut at different levels to generate 1, 2, …. *s* clusters, where *s* is the number of seizures. At each number of clusters *k*, the gap statistic *G*(*k*) was computed by comparing the within-cluster dispersion of the observed seizures and the reference datasets. The multiple reference datasets also allowed calculation of the standard error of the gap statistic at each *k*, *SE*(*k*). The optimal number of clusters was defined as the smallest number of clusters where *G*(*k*) ≥ *G*(*k*+1) − *SE*(*k*+1), which identifies the point at which increasing the number of clusters provides little improvement in the clustering of the data (77).

### Comparison to temporal distances

For each patient, we computed a “temporal distance matrix” containing the amount of time elapsed (measured in days) between the onset times of each pair of seizures. Spearman’s correlation was computed between the upper triangular elements of the seizure dissimilarity matrix and the temporal distance matrix of each patient. Since the distances in each matrix were not independent observations, the Mantel test (78) was used to determine the significance of each correlation. Briefly, the rows and columns of one matrix were randomly permuted 10,000 times. The correlation between the two sets of upper triangular elements was re-computed after each permutation, resulting in a distribution of correlation values that described the expected correlation if there were no relationship between seizure dissimilarities and temporal distances. The *p*-value of the association was then defined as the proportion of permuted correlation that were greater than or equal to the observed correlation. To correct for multiple comparisons, the Benjamini-Hochberg false discovery rate (FDR) correction (79) was applied to the set of *p*-values computed across all patients (31 total tests). The correlation was considered significant if the associated adjusted *p*-value was less than 0.05.

### Computing temporal correlation patterns

To quantify how seizure dynamics change over different timescales in each patient, Spearman’s correlation between seizure dissimilarities and temporal distances was computed only for seizure pairs with temporal distances less than or equal to timescale *T*. *T* was scanned from 0.25 days up to the patient’s largest temporal distance in steps of 0.25 days. A timescale was excluded from the analysis if less than seven pairs of seizures occurred within the given timescale or if no new seizure pairs were added when the timescale was increased. The resulting set of correlations across various timescales were referred to as “temporal correlation patterns.”

### Modelling seizure dissimilarities and temporal correlation patterns

To determine the underlying processes that could produce the observed temporal correlation patterns, changes in seizure dynamics were modelled using the functions

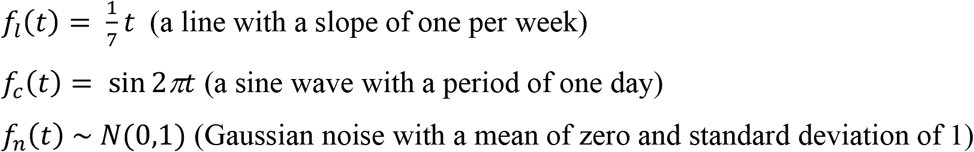

where *t* is time in days.

For each function, a simulated distance matrix *D* was then defined for the patients’ seizures, with

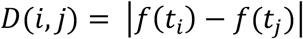

where *t*_*i*_ is the time of seizure *i*, *t*_*j*_ is the time of seizure *j*, and *f*(*t*) is the corresponding function. The dissimilarity of the two seizures was then defined as

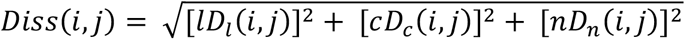

where *l*, *c*, and *n* are scalars controlling the relative contributions of the linear, circadian, and noise functions, respectively.

The relative contributions of the linear, circadian, and noise functions were scanned by varying the levels of *l*, *c*, and *n*. At each set of values, seizure dissimilarities were simulated 1000 times using different noise realisations (and correspondingly changing the noise distance matrix, *D*_*n*_), and the resulting temporal correlation patterns were computed for each set of simulated dissimilarities. Note that because temporal correlation patterns only depend on the order of the dissimilarities, only the relative magnitudes of *l*, *c*, and *n* affected the modelling results. A model was termed a “linear model” if *c* = 0, a “circadian model” if *l* = 0, and a “linear + circadian model” if *l* > 0 and *c* > 0.

To determine if a patient’s seizure dynamics could be categorised as linear, circadian, or linear + circadian, the simulated temporal correlation patterns were compared to the patient’s observed temporal correlation pattern by computing the mean squared error (MSE) of each simulated pattern. Simulated temporal correlation patterns with MSE ≤ 0.02185 were defined as “good matches” to the observed dynamics. This threshold was chosen because it was the 5^th^ percentile of the set of all MSEs, across all patients, and based on visual inspection of simulated temporal correlation patterns with different MSEs. The likelihood *L* of a given parameter set was then defined as the percentage of “good matches” produced by the 1000 noisy simulations of seizure dissimilarities at those parameter values. For each class of model (linear, circadian, or linear + circadian), the model’s likelihood (*L*_*l*_, *L*_*c*_, or *L*_*l+c*_, respectively) was the highest likelihood among the model type’s parameter sets, and the “best model” was the model with the highest likelihood. *L*_*n*_ was also defined as the highest likelihood of the parameter sets without any linear or circadian contributions (*l* = 0, *c* = 0, *n* > 0).

This best model with likelihood *L*_*max*_ was then used to categorise the patient’s dynamics if it outperformed all competing models. Specifically, we required that

1. The best model clearly outperform noise alone (*L*_*max*_ ≥ 2*L*_*n*_); otherwise, the patient’s dynamics were classified as other/indeterminate.
2. The performance of the linear model and circadian model were clearly distinguishable (*L*_*l*_ ≥ 2*L*_*c*_ if the linear model was best; *L*_*c*_ ≥ 2*L*_*l*_ if the circadian model was best); otherwise, the patient’s dynamics were classified as other/indeterminate.
3. If the best model was linear + circadian, it clearly outperform the two simpler models (*L*_*l+c*_ ≥ 2*L*_*l*_ and *L*_*l+c*_ ≥ 2*L*_*c*_); otherwise, the patient’s dynamics were classified as the simpler model (if one simpler model performed comparably by this criterion) or as other/indeterminate (if both simpler models performed comparably).

See SI Appendix, Text S8 for additional modelling details and the selected models for each patient.

## Code and data availability

All data was analysed using MATLAB version R2018b. To perform NMF, we used the Nonnegative Matrix Factorization Algorithms Toolbox, available at https://github.com/kimjingu/nonnegfac-matlab/, which implements the alternating nonnegative least squares with block principal pivoting algorithm (80, 81). For the remainder of the analysis, we used MATLAB implementations of standard algorithms (multidimensional scaling (Sammon mapping): mdscale (criterion “Sammon”), dynamic time warping: dtw, hierarchical clustering: linkage, Torgerson’s multidimensional scaling: cmdscale, gap statistic: evalclusters, FDR correction: mafdr) or custom code. The iEEG time series of all IEEG Portal patients is available at www.ieeg.org. The NMF factorisation of each patient’s data, along with the code for producing the primary downstream results (seizure dissimilarity matrices, clustering, and temporal analysis) and figures will be published on Zenodo (http://dx.doi.org/10.5281/zenodo.3560736).

## Acknowledgements

We thank Gerold Baier, Christoforos Papasavvas, Nishant Sinha, and the rest of the CNNP lab for discussions on the analysis and manuscript. We thank Andrew McEvoy and Anna Miserocchi for undertaking the epilepsy surgery at QS, and Catherine Scott, Roman Rodionov, and Sjoerd Vos for helping with data organisation.

The authors declare no conflict of interest.

## Author contributions

Conceptualization and methodology: GMS and YW. Investigation: BD. Resources: BD, PNT, and YW. Data curation: GMS, BD, PNT and YW. Software, formal analysis, visualization: GMS. Validation: YW. Project administration: GMS, PNT, and YW. Supervision, funding acquisition: PNT and YW. Writing – original draft preparation: GMS. Writing – review and editing: all authors.

## Supplementary information

Supplementary Information (Text S1-S11) is provided in the SI Appendix.

Patient-specific visualisations and results will be provided on Zenodo (http://dx.doi.org/10.5281/zenodo.3560736).

