## Supplementary material for "Seizure pathways change on circadian and slower timescales in individual patients with focal epilepsy": SI Appendix

- Text S1: Patient metadata
- Text S2: Dynamic time warping of example seizures
- Text S3: Summary of main analysis results of all patients
- Text S4: Amount of variability within and between seizure clusters
- Text S5: Seizure variability is not driven by differences in seizure ILAE clinical type
- Text S6: No relationship between features of seizure variability and clinical measures
- Text S7: No relationship between patterns of seizure dissimilarities and AED reduction
- Text S8: Supplementary modelling details and results
- Text S9: Hypothesised model for generating variability in seizure pathways
- Text S10: Dimensionality reduction using non-negative matrix factorization
- Text S11: Comparison of seizure dissimilarity to metric distances

### Text S1: Patient metadata

#### Table S1: Patient metadata

| patient | hospital | age (yrs) | sex | hemisphere | lobe | pathology | ILAE surgical outcome | total recording time | # seizures analysed | # electrodes analysed | sampling frequencies | AED reduction performed? |
|---|---|---|---|---|---|---|---|---|---|---|---|---|
| **Study 012-2** | MC | 37 | M | B | T | Other | - | 13d 16h | 28 | 81 | 499.907 Hz | - |
| **Study 017** | MC | 39 | M | R | FT | Other | 4 | 7d 17h | 9 | 16 | 499.907 Hz | - |
| **Study 019** | MC | 33 | M | L | T | - | 5 | 5d 16h | 33 | 96 | 499.907 Hz | - |
| **Study 020** | MC | 10 | M | R | F | - | 4 | 5d | 8 | 55 | 499.907 Hz | - |
| **Study 021** | MC | 16 | M | R | FT | Other | 1 | 6d 11h | 13 | 108 | 500 Hz | - |
| **Study 024** | MC | 23 | F | B | TP, IH | - | - | 8d 10h | 12 | 83 | 500 Hz | - |
| **Study 026** | MC | 9 | M | L | F | FCD | 1 | 3d 3h | 21 | 81 | 499.907 Hz | - |
| **Study 027** | MC | 34 | F | L | T | HS | - | 3d 21h | 6 | 47 | 500 Hz | - |
| **Study 030** | MC | 18 | F | L | FP | FCD | 3 | 5d 23h | 8 | 63 | 500 Hz | - |
| **Study 033** | MC | 3 | M | L | F | TS | 5 | 6d 17h | 17 | 127 | 500 Hz | - |
| **Study 037** | MC | 62 | F | R | F | - | - | 8d 23h | 8 | 78 | 499.907 Hz | - |
| **Study 038** | MC | 58 | M | L | FT | - | 1 | 3d | 10 | 86 | 500 Hz | - |
| **I002_P006_D01** | HUP | 26 | F | R | T | - | - | 12d 22h | 7 | 83 | 512 Hz | yes |
| **95** | UCLH | 35 | M | L | OP | Other | 4 | 7d 1h | 13 | 56 | 512 Hz, 1024 Hz | no |
| **756** | UCLH | 38 | F | B | T | Other | 3 | 6d 19h | 6 | 20 | 1024 Hz | yes |
| **770** | UCLH | 25 | F | L | P | FCD | 3 | 4d 3.6h | 8 | 71 | 512 Hz | no |
| **821** | UCLH | 25 | F | L | T | HS, BDI | 1 | 6d 4h | 9 | 46 | 512 Hz | yes |
| **931** | UCLH | 28 | M | L | T | HS | 4 | 7d | 11 | 58 | 512 Hz | yes |
| **934** | UCLH | 28 | F | R | OP | TS | 1 | 1d 19h | 40 | 76 | 512 Hz | no |
| **999** | UCLH | 28 | M | L | F | FCD | 1 | 12d 5h | 26 | 73 | 512 Hz | yes |
| **1005** | UCLH | 21 | F | R | T | HS | 2 | 8d 21h | 15 | 85 | 512 Hz | yes |
| **1097** | UCLH | 28 | M | L | F | GL | 1 | 1d 20h | 8 | 84 | 512 Hz | no |
| **1109** | UCLH | 31 | F | R | T | CAV | 1 | 6d 1h | 13 | 53 | 1024 Hz | yes |
| **1149** | UCLH | 43 | F | R | TOP | DNT | 1 | 7d 22h | 24 | 62 | 512 Hz, 1024 Hz | no |
| **1163** | UCLH | 27 | F | L | F | FCD | 1 | 8d | 8 | 111 | 512 Hz | yes |
| **1167** | UCLH | 39 | M | L | P | CAV | 4 | 5d 22h | 43 | 51 | 1024 Hz | no |
| **1168** | UCLH | 60 | F | L | F | FCD | 2 | 2d | 10 | 94 | 512 Hz | no |
| **1182** | UCLH | 28 | M | R | P | FCD | 3 | 5d 5h | 52 | 75 | 512 Hz | no |
| **1196** | UCLH | 41 | M | R | T | HS | 3 | 15d 22h | 11 | 34 | 1024 Hz | yes |
| **1200** | UCLH | 24 | F | R | T | HS | 1 | 2d 19h | 14 | 71 | 512 Hz | yes |
| **1211** | UCLH | 26 | M | R | T | Other | 3 | 5d 7h | 20 | 77 | 512 Hz | yes |

Table S1 lists the metadata for the patients whose seizures were analysed in this study. Patient identifiers are listed under "patient." For IEEG Portal patients (MC and HUP hospitals), their identifier is the same as the one used by the database. Metadata was extracted from the reports provided on the IEEG Portal (MC and HUP patients) or the patient clinical reports (UCLH patients). For each patient, the following information is provided:

- hospital: hospital at which the patient underwent presurgical monitoring (MC = Mayo Clinic, HUP = Hospital of the University of Pennsylvania, UCLH = University College London Hospital).
- age: age, in years, at the time of the presurgical monitoring.
- sex: patient sex (M = male, F = female).
- hemisphere: purported hemisphere of onset of the patient's seizures (L = left, R = right, B = bilateral), based on clinical findings.
- lobe: purported lobe of onset of the patient's seizures (T = temporal, F = frontal, P = parietal, O = occipital, IH = interhemispheric), based on clinical findings. Note that some patients had seizures arising from multiple lobes/at the boundary of two lobes (e.g., OP = occipital/parietal onset).
- pathology: postoperative tissue pathology findings (FCD = Focal cortical dysplasia, BDI = Brain damage - inflammatory, HS = Hippocampal sclerosis, TS= Tuberous sclerosis, GL = Glioma, CAV = Cavernoma, DNT = Dysembryoplastic neuroepithelial tumour, Other = other type of pathology that is not one of the other categories). A dash indicates that this information is unavailable.
- ILAE surgical outcome: patient surgical outcome according to the International League Against Epilepsy classification (1 = seizure free, 2 = only auras, 3+ = not seizure free). A dash indicates that the patient did not undergo surgery or their surgical outcome is unavailable. For IEEG Portal patients (MC and HUP hospitals), the surgical outcome provided by the database is given. For UCLH patients, the 12 months post-surgical outcome is provided.

- total recording time: total duration of the presurgical intracranial recording time.
- # seizures analysed: number of the patient's seizures analysed in this work.
- # electrodes analysed: number of recording electrodes included in the analysis, after removing noisy electrodes.
- sampling frequencies: sampling frequencies at which intracranial data was acquired and stored.
- AED reduction performed: whether patient antiepileptic drugs (AEDs) were systematically reduced during the presurgical recording. A dash indicates that this information is unavailable.

### Text S2: Dynamic time warping of example seizures

In this section, we demonstrate how dynamic time warping aligns similar dynamics across seizures. In our application, this warping allows us to identify seizures with similar network evolutions, even if the rates of the evolutions differ.

As a reminder, dynamic time warping stretches a pair of two time series in order to minimise the distance between them. Importantly, dynamic time warping can *only* stretch each time series in order to align similar dynamics, which means that the algorithm cannot

1) skip time points; instead, all time points of both time series, including beginning points, must be included in the warp path.
2) repeat earlier time points; i.e., the warp path cannot double back on itself in order to repeatedly include a section of one of the time series.

The warping process is also repeated individually for each pair of seizures, and the warp length and path will therefore differ between different pairs of seizures. In order words, this algorithm provides a *pairwise* alignment of time series, rather than a multiple time series alignment. We focus on pairwise alignments because a global warping solution would likely sacrifice the optimal alignment of some pairs of time series.

To understand how dynamic time warping aligns pairs of time series, consider two seizures, seizure A with M windows, and seizure B with N windows. From these two seizures, we compute the time-time distance matrix D (Fig. S2a). This MxN matrix contains the pairwise distance between the functional connectivity of each pair of time windows across the two seizures: D(m,n) is the distance between the functional connectivity of the m-th time window of seizure A and the n-th time window of seizure B. Thus, the row of D corresponds to the time window index of seizure A, while the column of D corresponds to the time window index of seizure B.

Dynamic time warping finds a path through this distance matrix that minimises the total distance (here, the L1 norm distance) between the two seizures. The algorithm only allows three categories of moves through the matrix (Fig. S2a):

1) Horizontal moves, D(m,n) → D(m,n+1), stretch seizure A (blue arrow)
2) Vertical moves, D(m,n) → D(m+1,n), stretch seizure B (red arrow)
3) Diagonal moves, D(m,n) → D(m+1,n+1), do not stretch either seizure (purple arrow)

We demonstrate this process by visualising the time-time distance matrices and warp paths of three pairs of patient 1109's seizures (Fig. S2b-d). Each time-time distance matrix is computed by calculating the L1 norm distance between the functional connectivity vectors (reconstructed following NMF – see Supplementary Fig. S10.2) of each pair of time windows in the two seizures. Low distances reveal pairs of time windows with similar functional network dynamics. Points in the warp path (red squares), laid over the time-time distance matrix, indicate which pairs of indices were aligned. Finally, to calculate the seizure dissimilarity between two seizures, we average over the pointwise distances along the warp path. To visually demonstrate how DTW aligns similar time windows, we additionally assigned each seizure time window to a network state (see Supplementary Fig. S10.2); each seizure can therefore be described as a progression of different network states. Time windows with the same state have similar functional network connectivity and should therefore, when possible, be aligned across different seizures.

To provide a simple example of the warping process, we first examine the warp path of seizures 7 and 12, which have a relatively low seizure dissimilarity (0.62) (Fig. S2b). Both seizures begin with network state 3 (orange) and then transition to network state 1 (aqua). While the duration of state

3 is the same in both seizures, state 1 lasts longer in seizure 12 than seizure 7. As such, window 7 of seizure 7 is repeated in order to align the two seizures, as shown by the horizontal line in the warp path.

Fig. S2c shows the warp path of seizures 3 and 4, which have similar state progressions but differ in the rate of their state progressions. However, because dynamic time warping aligns the similar parts of each seizure, there is also a relatively low seizure dissimilarity between these two seizures (0.52). These two seizures exemplify how certain windows are stretched to accommodate longer state durations in the other seizure: horizontal lines in the path correspond to places where seizure 3 is stretched, while vertical lines correspond to time points in seizure 4 that are repeated. For example, in seizure 3, a window of the last state (state 5, green) is stretched so that it matches the dynamics of state 4. Notably, the brief transition of seizure 4 to state 6 (dark orange) cannot be matched by any time point in seizure 3 because the warp path cannot skip this window or align it to earlier windows. In seizure 4, the longest warpings occur during state 3 (orange) and state 2 (wine red), which are both longer in seizure 3. Thus, by warping the seizure time series, our method recognises parts of the seizures that have similar network dynamics, despite differences in the rates of the seizure evolutions.

Finally, Fig. S2d demonstrates how dynamic time warping aligns two seizures, seizures 3 and 6, that have somewhat different state progressions. Compared to seizure 3, seizure 6 spends more time in state 2 (wine red) and never progresses to seizure 3's final state, state 5 (green). As such, in seizure 3, a window of state 2 is stretched. However, because the warp path must include all time points, the final part of seizure 6 must be matched to the final dynamics of seizure 3, even though there are high distances between these time points. This difference between the final seizure dynamics raises the dissimilarity between seizures 3 and 6: it is 1.03, which is almost double the previously examined dissimilarity between seizures 3 and 4.

Note that seizures 3 and 6 provide an example of when seizure pathways only partially match: the dynamics are initially very similar, and seizure 3 diverges by progressing to an additional state. In these cases, dynamic time warping cannot find a close alignment between the entire seizure progressions, and our seizure dissimilarity measure depends on the durational proportion of the mismatch. In other words, had the green state lasted longer in seizure 3, the seizure dissimilarity measure would be higher in this example. In some cases, this dependence on state duration also means that the seizure dissimilarity measure is not a metric distance. In Supplementary S11 we compare seizure dissimilarities, computed using dynamic time warping, to two alternative metric distance measures for comparing seizure pathways.

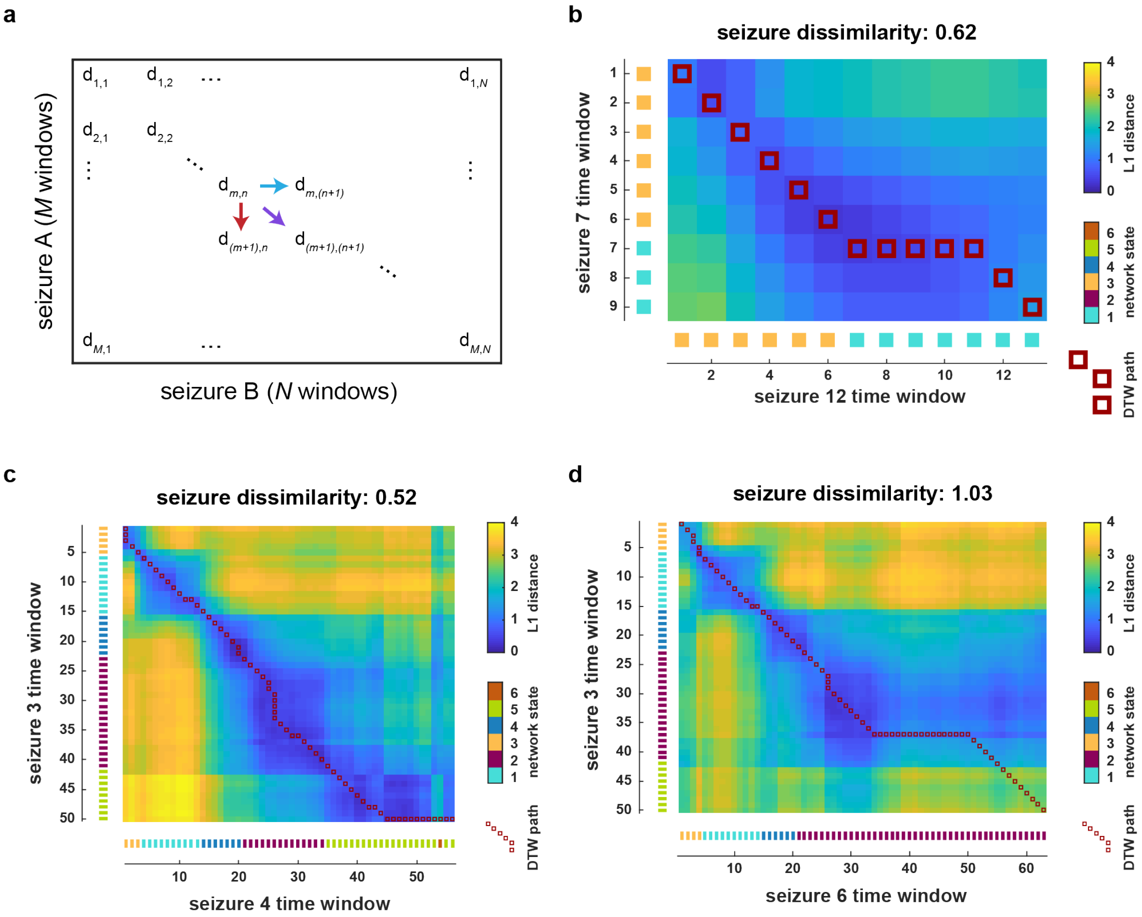


**Fig. S2: Visualising the optimal path found by dynamic time warping.** (a) The time-time distance matrix, $D_{MxN}$, contains the distance between each pair of time windows across two seizures. A warp path must start at D(1,1) and move horizontally, vertically, or diagonally until it reaches D(M,N). (b-d) The warp path and time-time distance matrices of patient 1109's (b) seizures 7 and 12, (c) seizures 3 and 4, and (d) seizures 3 and 6. Each heatmap shows the L1 distance between the functional network evolutions of the pair of seizures. Low distances, in blue, indicate time points with similar network dynamics. Red squares indicate the warp path chosen by the dynamic time warping algorithm; if entry (i,j) is part of the warp path, then the corresponding time windows of the two seizures were aligned. For example, in (b), seizure 7's time window 7 is aligned to seizure 12's time windows 7-11 . Along each side of the time-time distance matrix, the network state of each time point is shown. Note that since the time-time distance matrix is calculated from the functional connectivity time courses, and not the simplified state descriptions, there are non-zero distances between points assigned to the same state. However, the network state progressions provide a useful visualisation for how the warp path aligns comparable network states.

### Text S3: Summary of main analysis results of all patients

### Table S3: Summary of main analysis results of all patients

| patient | number of seizures | number of clusters | number of states | correlation with temporal distance | | | Model matching temporal correlation pattern |
|---|---|---|---|---|---|---|---|
| | | | | *rho* | *p-value* | *q-value* | |
| **Study 012-2** | 28 | 2 | 8 | 0.57 | <0.0001 | <0.0002 | Linear |
| **Study 017** | 9 | 1 | 4 | 0.28 | 0.0990 | 0.1228 | Other/indeterminate |
| **Study 019** | 33 | 1 | 9 | 0.43 | <0.0001 | <0.0002 | Linear |
| **Study 020** | 8 | 1 | 7 | 0.48 | 0.0118 | 0.0205 | Linear |
| **Study 021** | 13 | 1 | 5 | 0.67 | 0.0001 | 0.0002 | Linear |
| **Study 024** | 12 | 2 | 6 | 0.67 | <0.0001 | <0.0002 | Linear |
| **Study 026** | 21 | 1 | 4 | 0.53 | <0.0001 | <0.0002 | Linear |
| **Study 027** | 6 | 1 | 7 | 0.56 | 0.0119 | 0.0205 | Linear + circadian |
| **Study 030** | 8 | 2 | 8 | 0.72 | 0.0216 | 0.0319 | Linear |
| **Study 033** | 17 | 1 | 4 | -0.10 | 0.8567 | 0.8567 | Other/indeterminate |
| **Study 037** | 8 | 1 | 10 | 0.58 | 0.0161 | 0.0263 | Linear |
| **Study 038** | 10 | 1 | 8 | 0.19 | 0.1572 | 0.1805 | Other/indeterminate |
| **I002_P006_D01** | 7 | 1 | 7 | 0.83 | 0.0035 | 0.0072 | Linear + circadian |
| **95** | 13 | 2 | 3 | 0.71 | <0.0001 | <0.0002 | Linear |
| **756** | 6 | 2 | 6 | 0.18 | 0.1724 | 0.1909 | Linear |
| **770** | 8 | 1 | 5 | 0.09 | 0.2598 | 0.2685 | Circadian |
| **821** | 9 | 2 | 4 | 0.17 | 0.1304 | 0.1555 | Linear + circadian |
| **931** | 11 | 1 | 5 | 0.69 | 0.0001 | 0.0002 | Linear |
| **934** | 40 | 2 | 3 | 0.63 | <0.0001 | <0.0002 | Linear |
| **999** | 26 | 1 | 5 | 0.64 | <0.0001 | <0.0002 | Linear |
| **1005** | 15 | 1 | 4 | 0.69 | <0.0001 | <0.0002 | Linear + circadian |
| **1097** | 8 | 1 | 3 | 0.40 | 0.0494 | 0.0666 | Linear |
| **1109** | 13 | 3 | 6 | 0.24 | 0.0527 | 0.0681 | Linear |
| **1149** | 24 | 1 | 8 | 0.42 | 0.0001 | 0.0002 | Linear |
| **1163** | 8 | 1 | 8 | 0.12 | 0.2559 | 0.2685 | Other/indeterminate |
| **1167** | 43 | 2 | 5 | 0.22 | 0.0115 | 0.0205 | Circadian |
| **1168** | 10 | 2 | 2 | 0.31 | 0.0363 | 0.0512 | Linear + circadian |
| **1182** | 52 | 2 | 6 | 0.65 | <0.0001 | <0.0002 | Linear + circadian |
| **1196** | 11 | 1 | 5 | 0.32 | 0.0208 | 0.0319 | Linear + circadian |
| **1200** | 14 | 1 | 4 | 0.69 | <0.0001 | <0.0002 | Linear |
| **1211** | 20 | 1 | 4 | 0.38 | 0.0003 | 0.0007 | Circadian |

Table S3 summarises the main analysis results of all patients. See main text Methods for a detailed description of the analysis. Patient identifiers are listed under "patient." The next columns of the table provide the following information for each patient:

- number of seizures: the number of seizures analysed.
- number of clusters: the number of non-hierarchical seizure clusters, based on the seizure dissimilarity matrix.
- number of states: the optimal number of network states (i.e., NMF basis vectors), which was used to reduce noise in the dataset. See Supplementary Fig. S10.1 for details on finding the optimal number of states.

The table then provides the following information about the correlation between seizure dissimilarities and temporal distances (amount of time elapsed between seizure start times) in each patient:

- rho: the Spearman correlation between seizure dissimilarities and temporal distances
- *p*-value: the *p*-value of the correlation, based on permutation tests (10,000 permutations). If a value greater than or equal to the observed correlation was not observed, then the *p*-value is listed as < 0.0001.

- *q*-value: the *q*-value, or adjusted *p*-value, of the correlation after global false discovery rate correction on all *p*-values in the table (31 total statistical tests).

Significant correlations (defined as $q < 0.05$) are indicated with blue text for the correlation and the corresponding *p*- and *q*-values.

The final column in the table ("Model matching temporal correlation pattern") indicates the type of model consistent with each patient's temporal correlation pattern (see "Seizure pathways change on different timescales" in main text). Detailed modelling results are provided in Supplementary S8.

Visualisations of the analysis results of each patient will be available on Zenodo at http://dx.doi.org/10.5281/zenodo.3560736.

**Text S4: Amount of variability within and between seizure clusters**

In each patient, seizures were clustered based on seizure dissimilarities (see main text Methods, "Seizure clustering and cluster evaluation"). Each patient either had a spectrum of seizures (one seizure cluster, with variability in seizure pathways within that cluster) or two or more seizure clusters (i.e., their seizure pathways could be grouped into different types of dynamics, with more similarity within a group than between groups). Fig. S4 shows the median level of seizure dissimilarity of seizures within the same cluster and the median level of seizure dissimilarity of seizures from different clusters in each patient. Overall, in patients with multiple seizure clusters (purple histograms), the average seizure dissimilarity *between* clusters is higher than the average seizure dissimilarity *within* clusters, as expected. However, there is overlap in the distributions of within- and between-cluster seizure dissimilarity, demonstrating that seizures in the same cluster can be relatively different, while seizures in different clusters can be relatively similar. In other words, a given cluster can represent *different* pathways that are, despite their diversity, still more similar to each other than to the patient's other seizure pathways. Meanwhile, in some patients, the different types of seizure pathways, represented by different clusters, are still relatively similar, indicating overall lower variability in seizure dynamics in that patient.

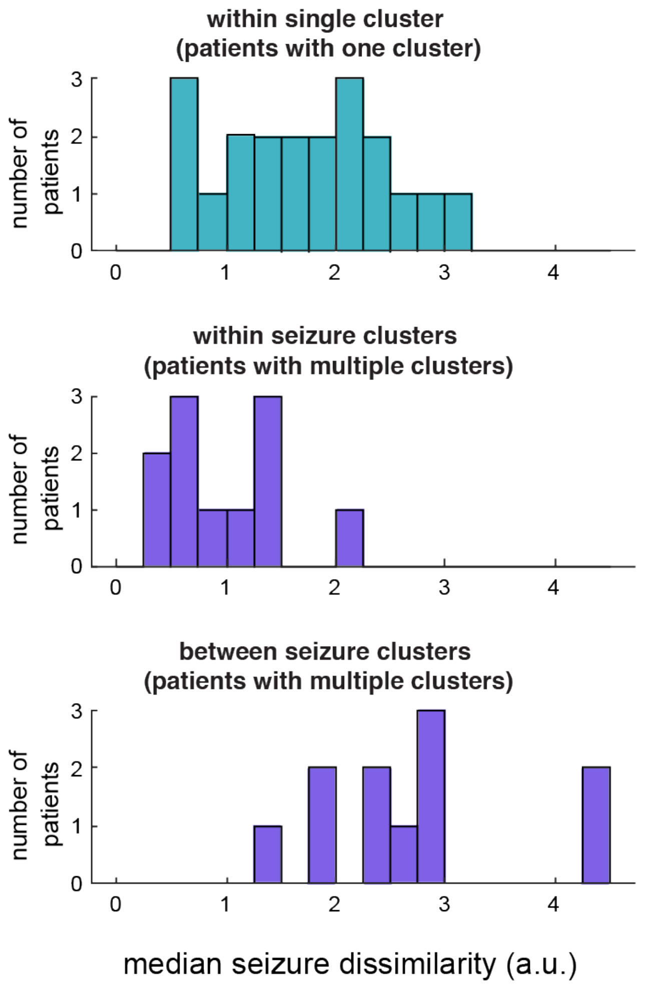


**Fig. S4: Amount of variability within and between seizure clusters.** Teal correspond to data from patients with a single seizure cluster, while purple corresponds to data from patients with two or more seizure clusters. Top histogram: median within-cluster seizure dissimilarity in patients with one seizure cluster. Note that since all seizures are in the same cluster in these patients, these values are the same as the overall median seizure dissimilarity of these patients. Thus, this histogram reproduces the teal histogram shown in Fig. 2C of the main text. Middle histogram: median within-cluster seizure dissimilarity in patients with multiple seizures clusters. Bottom histogram: median between-cluster seizure dissimilarity in patients with multiple seizure clusters.

**Text S5: Seizure variability is not driven by differences in seizure ILAE clinical type**

In this section, we qualitatively examine whether our quantitative comparison of within-patient seizure network dynamics provides more information about seizure dynamics than the International League Against Epilepsy (ILAE) clinical seizure classification alone(15–17). Importantly, unlike our seizure dissimilarity measure, the ILAE clinical seizure classification was not designed to group seizures based on their dynamics; rather, a type is defined as "a useful grouping of seizure characteristics for purposes of communication in clinical care, teaching, and research(17)." Nonetheless, comparing the seizure groupings based on dynamics and clinical classification reveals if the observed variability is solely explained by differences in seizure clinical classification; for example, it is possible that the observed variability could be solely attributed to focal seizures that all have similar initial dynamics, but sometimes secondarily generalise.

Based on the clinical reports for each patient, seizures were labelled as

- subclinical: seizures that are visible electrographically, but do not cause any apparent symptoms
- focal: seizures that originate in networks within one hemisphere; here, we limit this label to seizures that also remain focal (i.e., do not secondarily generalise)
- secondarily generalised: seizures that begin focally and subsequently engage bilateral networks, resulting in convulsions that have tonic and/or clonic components

Note that secondarily generalised seizures now correspond to "focal to bilateral tonic clonic seizures" in the more recent ILAE clinical classification(17); however, to be consistent with the terminology in the patients' reports and the previous literature, we use the older classification term here. Additionally, focal seizures can be further subdivided into different categories based on, for example, whether awareness is preserved during the seizure. However, we do not make those designation here due to the absence of this information or uncertainty in the classification in the many of the clinical reports.

The dendrograms of Fig. S5 show the hierarchical clustering of seizures, based on seizure network dynamics (i.e., based on our seizure dissimilarity measure) in example patients. More similar seizures, represented by leaves on the dendrogram, are joined by nodes. The height of the node linking two seizures (or groups of seizures) represents the dissimilarity between them, with higher nodes indicating less similar seizures. The seizures are additionally labelled by their ILAE clinical type (subclinical, focal, or secondarily generalised) to examine the relationships between seizures of the same and different clinical type(s).

In some patients, only a single clinical seizure type was available/suitable for analysis. Fig. S7a provides examples of three patients in which only focal seizures were analysed. Although all seizures shared the same clinical type, there was variability in the seizure network dynamics in each patient. For example, based on the dendrogram of patient 1005, seizures 13, 14, and 15 appeared to have similar dynamics, but overall this group was different from the remaining seizures. These patients alone demonstrate that the observed variability is not solely due to differences in clinical seizure type, as there is variability within a single clinical type. The variability within focal seizures is unsurprising, as focal seizures can have varying levels of spread and symptoms, indicating diversity in their spatial spreads and severity. Our hierarchical clustering based on dynamics also suggests that focal seizures can also be associated with different patterns of brain network interactions.

In other patients, we saw that, based on their network dynamics, seizures could be divided into groups that correspond to their clinical types (Fig. S5b). In patients 821 and 1163, the dendrograms can be cut such that the resulting clusters match the ILAE clinical classification: each cluster only contains one clinical type, and each cluster also contains all examples of the clinical type. However, note that this grouping of the seizures is not necessarily the *optimal* non-hierarchical clustering of the seizures (see main text Methods, "Seizure clustering and cluster evaluation," for how optimal clusters were determined using the gap statistic). Indeed, even in these patients, we observed that high levels of variability within some clinical types. For example, in patient 1163, there were large amounts of variability within the focal seizures. Thus, even in the patients with close agreement between clinical types and dynamical clusters, there was additional variability in seizure network dynamics that was not explained by the coarse clinical classification alone.

Finally, in other patients, we observed that the seizures' clinical classifications do not perfectly align with the hierarchical clustering based on the seizures' network dynamics (Fig. S5c). In these patients, cutting the dendrograms at different levels produced clusters that contain multiple clinical types and/or more than one cluster containing a given clinical type. For example, partitioning patient I002_P006_D01's seizures to produce four clusters would segregate the secondarily generalised seizures and subclinical seizures, as these types have fairly homogeneous dynamics in the observed seizures. However, the two focal seizures, 3 and 7, would form separate groups due to their disparate dynamics. Meanwhile, in other patients, we saw that seizures can be more similar to seizures of different clinical types than seizures of the same clinical type. For example, in patient 1200, seizure 4 (focal) was grouped with subclinical seizures, seizures 5, 6, and 7, while seizure 9 (also focal) appeared to be very similar to a different subclinical seizure, seizure 11. Similarities across clinical types is also unsurprising, as seizures of different clinical types can share similar properties, such as ictal rhythms and propagation pathways. Our analysis suggests that similar brain network interactions can also occur in seizures of different clinical types. Notably, the similarities between focal and subclinical seizures also suggests that seizures can share common network features even when their symptoms differ.

In summary, although seizures of the same clinical type often shared similar network dynamics, we also observed that, in a given patient, 1) there was variability in seizure network evolutions within seizures of the same clinical type, and 2) seizures of different clinical types could share aspects of their network dynamics. These results are unsurprising given that the coarse clinical classification used here is not designed to group seizures based on their network dynamics. Indeed, more generally, seizures of the same clinical type are known to have different features, while seizures of different clinical types can have similar dynamics.

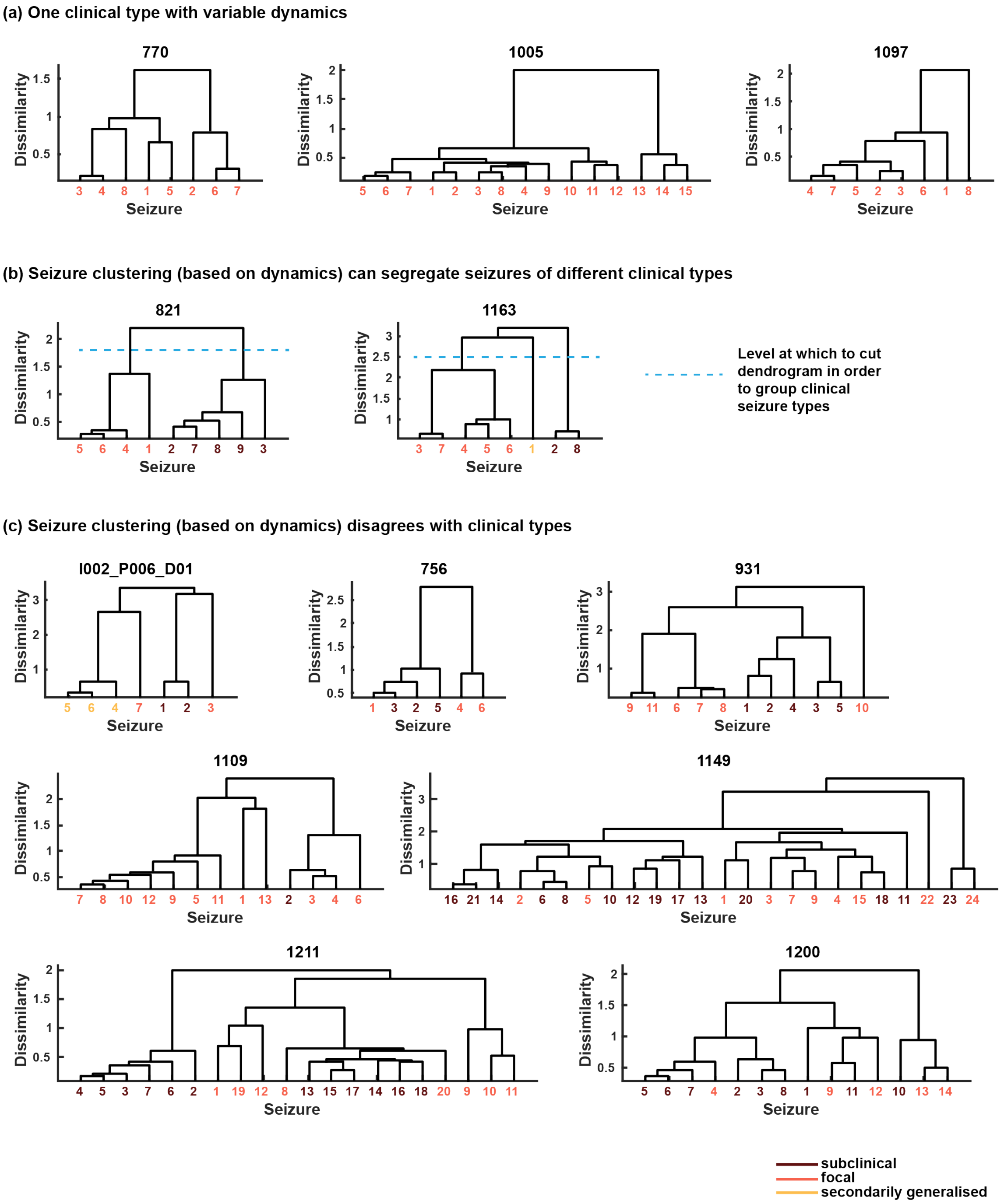


**Fig. S5: Comparison of ILAE clinical seizure types classification and variability in seizure network dynamics.** (a-c) Dendrograms of example patients with known clinical classifications for all seizures. Each dendrogram describes the hierarchical clustering of the patient's seizures, based on the seizure network evolutions (see Methods of main text). More similar seizures, represented by leaves on the dendrogram, are joined by nodes. The height of the node linking two seizures (or groups of seizures) represents the dissimilarity between them, with higher nodes indicating less similar seizures. Seizure labels are coloured by their ILAE clinical type (dark red = subclinical, orange = focal, yellow = secondarily generalised). (a) Example patients whose analysed seizures consisted of a single clinical type (focal). Note that there was variability in seizure network dynamics in each patient. (b) Example patients whose analysed seizures consisted of at least two clinical types, and whose hierarchical clustering of seizures agreed with the clinical seizure classification; i.e., the dendrogram can be cut at a specific level (blue dotted line) to perfectly segregate seizures of different clinical types. (c) Example patients whose hierarchical clustering did not agree with the seizure clinical classification; i.e., there is no way to cut the dendrogram to perfectly segregate seizures of different clinical types. The resulting clusters will contain multiple clinical types and/or multiple clusters will contain the same clinical type.

**Text S6: No relationship between features of seizure variability and clinical measures**

In this section, we explore if certain features of seizure variability are associated with clinical features, such as seizure onset location or surgical outcome.

***Comparison of clinical features to temporal distance correlation, number of clusters, and median seizure dissimilarity***

For each patient, we first computed three different features that describe aspects of seizure variability:

1. Temporal distance correlation: the correlation between seizure dissimilarities and temporal distances
2. Number of clusters: the optimal number of seizures clusters, based on seizure network dynamics (i.e., based on the seizure dissimilarity matrix)
3. Median seizure dissimilarity: the average seizure dissimilarity across all pairs of seizures (computed by taking the median of the upper triangular elements of the seizure dissimilarity matrix)

See Methods of the main text for details on computing these measures.

For each measure, we compared

- Patients labelled as having strictly temporal lobe onset seizures (n = 12) vs. strictly frontal lobe onset seizures (n = 8)
- Patients labelled as having left hemisphere onset seizures (n = 15) vs. right hemisphere onset seizures (n = 13)
- Male patients (n = 16) vs female patients (n = 15)

We were unable to compare seizure variability features in bilateral onset patients or in patients with certain pathological features (e.g., hippocampal sclerosis) due to small sample sizes.

For each group and feature, we first used the Kolmogorov Smirnov test to determine if the group's feature distribution was consistent with a normal distribution. If so, we also used a two-sample F-test to test the null hypothesis that the feature distributions of two groups being compared (e.g., temporal and frontal patients) had equal variances. For these pairs of distributions, we failed to reject the null hypothesis in all cases, allowing us to assume equal variances and compare the group means using a two-tailed Student's *t*-test. If, however, the normality assumption was not satisfied, a Wilcoxon rank sum test was instead used to compare the group medians.

Additionally, we used Spearman's correlation to evaluate the relationship between the surgical outcome (scored according to ILAE criteria) and each of the three features in the 26 patients who underwent surgical resection and had a known surgical outcome (see Table S1 for the ILAE surgical outcome of each patient). An ILAE score of 1 indicates complete seizure freedom after surgery, with successively higher scores indicating worse outcomes.

For all features, we found no significant differences (defined as a $p$-value < 0.05) between temporal and frontal lobe patients, left and right hemisphere onset patients, or male and female patients (Fig. S6A-C). Additionally, there was no significant relationship between ILAE surgical outcome and any of the seizure variability features (Fig. S6D). Our results suggest that these clinical features do not impact the amount or form of seizure variability, or that any effect is smaller than detectable with our sample sizes. Further research is needed to determine the factors that shape features of seizure variability.

#### *Comparison to model timescales (see main text, "Seizure pathways change on different timescales")*

We additionally investigated whether ILAE surgical outcome differed between patients with different categories of seizure variability (linear, circadian, or linear + circadian). For the 22 patients with both an assigned model category (i.e., their model category was not "other/indeterminate") and a known ILAE surgical outcome, we used ordinal regression to test for a significant relationship between ILAE surgical outcome and model type. The two independent variables were whether the patient's model had a linear component (i.e., if the patient belonged to either the linear or linear+circadian category) and whether the patient's model had a circadian component (i.e., if the patient belonged to either the circadian or linear+circadian category). There was no significant relationship between these variables and ILAE surgical outcome.

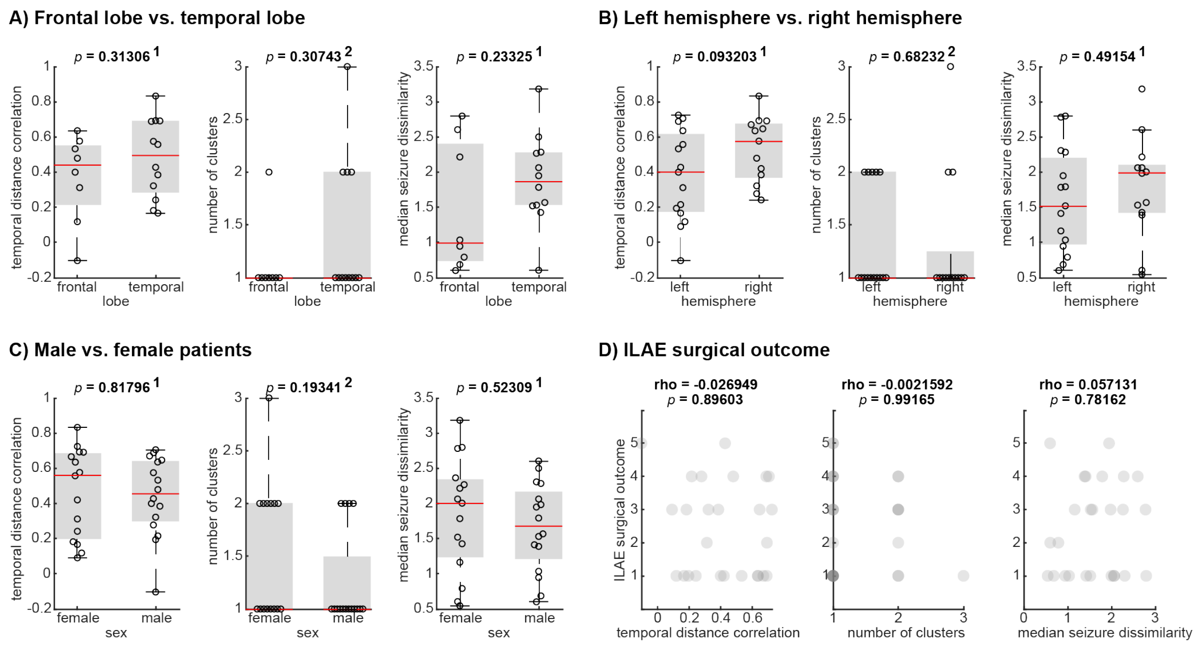


**Fig. S6: Comparison of seizure variability features in different groups of focal epilepsy patients.** A) Comparison of features in patients with frontal lobe onset vs. patients with temporal lobe onset. B) Comparison of features in patients with left hemisphere onset vs. right hemisphere onset. C) Comparison of features in male vs. female patients. D) Features vs. patient ILAE surgical outcome. In A, B, and C, each subplot shows the distribution of a seizure variability feature in the corresponding patient subgroups. See the text of Supplementary S6 for a description of each feature. Red horizontal lines mark the median feature value for each population, while the lower and upper bounds of each grey box indicate the first and third quartiles, respectively, of each distribution. For each feature, an appropriate test (superscript 1 = two-tailed Student's t-test, 2 = Wilcoxon rank sum test; see Supplementary S6 text for details) was used to compare group means or medians, yielding the given *p*-values. In D, each scatterplot shows the relationship between a seizure variability feature and patient ILAE surgical outcome. Note that in some plots, points overlap, especially when the seizure variability feature only takes on discrete values (e.g., the number of seizure clusters); in these cases, the darkness of the point indicates the number of patients it represents. The association between surgical outcome and each feature was quantified using Spearman's correlation, and the resulting correlation rho and associated *p*-value is given above each scatter plot.

### S7: No relationship between temporal patterns of seizure dissimilarities and AED reduction

#### *Comparison to overall correlation with temporal distances (see main text, "Seizures with more similar pathways tend to occur closer together in time")*

During presurgical recordings, the antiepileptic drug (AED) dosages of patients are often gradually reduced to provoke more seizures, and thus improve localization of the epileptogenic zone(9–12). Because such medication alters neural excitability(13, 14), it is possible that it also affects seizure dynamics. Thus, gradual changes in AED dosages could potentially produce gradual changes in seizure dynamics, leading to the observed temporal changes in seizure dynamics in many patients. We therefore investigated if differences in the correlation between seizure dissimilarities and temporal distances across patients could be attributed to whether antiepileptic medications were reduced in each patient.

Information about medication dosages was available from the clinical reports of nineteen patients. This information was extracted and used to label each patient as "AED reduction performed" or "no AED reduction performed." Medication changes due to stat doses of medication (i.e., medication given in addition to the planned dose in order to control seizures) were not considered in this assessment; rather, we sought to identify whether gradual changes in medication were intentionally made, as those could underlie the observed gradual temporal changes in seizure dynamics. We then also labelled whether each of the 19 patients had a significant or not significant correlation between seizure dissimilarities and temporal distances.

Table S7.1 shows the cross-tabulation table of these variables. If AED reduction reliably altered seizure dynamics, we would expect significant correlations between seizure dissimilarities and temporal distances in approximately all patients who underwent medication reduction. However, of the eleven patients that underwent AED reduction, four did *not* have a significant association between these distances. Additionally, if the temporal changes in seizure dynamics were *solely* due to medication reduction, we would not expect a significant relationship between seizure dissimilarities and temporal distances in patients who did *not* undergo AED reduction. Instead, we observed a significant correlation in five of the eight patients who did not undergo AED reduction. A $\chi^2$ test found no association between whether AED reduction was performed and whether the correlation between seizure dissimilarities and temporal distances was significant ($\chi^2 = 0.0026$, $p = 0.96$). As such, the temporal relationship between similar seizure dynamics cannot solely be attributed to AED reduction in our cohort.

**Table S7.1: Cross-tabulation table describing the frequency of antiepileptic drug (AED) reduction (columns, "AED reduction performed") and the frequency of a significant correlation between temporal distances and seizure dissimilarities (rows, "Significant correlation") in our cohort of patients.** For example, three patients did not undergo AED reduction and did not have a significant correlation between temporal distance and seizure dissimilarities (AED reduction performed: no, significant correlation: no). At the end of each row and column, the total count of patients in the respective category is also given (e.g., a total of 7 patients did not have a significant correlation between temporal distance and seizure dissimilarities).

| | | AED reduction performed | | |
|---|---|---|---|---|
| | | **no** | **yes** | **total** |
| **Significant correlation** | **no** | 3 | 4 | 7 |
| | **yes** | 5 | 7 | 12 |
| | **total** | 8 | 11 | **19** |

#### *Comparison to model timescales (see main text, "Seizure pathways change on different timescales")*

We additionally explored if patients who underwent AED reduction had seizure dynamics best described by a particular model of seizure variability (linear, circadian, or linear + circadian dynamics). As a reminder, the linear model describes more gradual changes in seizure dynamics over the course of the recording, which could be attributed to AED reduction.

Table S7.2 shows the number of patients with and without AED reduction who were assigned to each model category. A $\chi^2$ test cannot be performed here due to small sample sizes and assignment to multiple categories (patients with linear + circadian variability could be considered members of both the linear and circadian groups). However, from the cross-tabulation table alone, it is apparent that similar proportions of patients with and without AED reduction were assigned to each model category. In particular, AED reduction alone cannot explain the linear pattern of seizure variability: six of the eight patients without AED reduction had variability that was categorised as either linear or linear + circadian. Thus, factors beyond AED reduction influence the temporal patterns of changes in seizure pathways in this cohort.

**Table S7.2: Cross-tabulation table describing the frequency (number of patients) of each model category (linear, circadian, linear + circadian, and other/indeterminate) among patients with and without AED reduction.**

| | **Linear** | **Circadian** | **Linear + circadian** | **Other/indeterminate** |
|---|---|---|---|---|
| **AED reduction** | 5 | 1 | 4 | 1 |
| **No AED reduction** | 4 | 2 | 2 | 0 |

### Text S8: Supplementary modelling details and results

#### *Model parameter scan*

**Table S8: Scan values for model parameters**

| Parameter | Description | Scan range | Scan step size |
|---|---|---|---|
| $l$ | Scales linear contribution (i.e., slow/gradual changes) | 0 to 1 | 1 |
| $c$ | Scales circadian (sinusoidal) contribution | 0 to 5 | 0.1 |
| $n$ | Scales noise contribution | 0.025 to 0.5 | 0.025 |

Table S8 shows the values used to scan the parameters $l$, $c$, and $n$ in the model of seizure temporal correlation patterns. As discussed in the main text Methods (see "Modelling seizure dissimilarities and temporal correlation patterns"), the parameters $l$, $c$, and $n$ control the relative contributions of the linear, circadian, and noise functions to the simulated temporal changes in seizure dynamics.

Note that temporal correlation patterns (see main text Methods, "Computing temporal correlation patterns") only depend on the relative magnitude of seizure dissimilarities, and not their absolute magnitude. This situation arises because only the order of dissimilarities affects Spearman's correlation between the seizure dissimilarities and temporal distances. Thus, since model parameters were selected based on how well they reproduced each patient's observed temporal correlation patterns, only the relative values of the model parameters matter for model selection. For example, given the same noise realisation, the temporal correlation patterns of the parameter sets ($l$ = 0.5, $c$ = 1, $n$ = 0.05) and ($l$ = 1, $c$ = 2, $n$ = 0.1) would be equivalent. These properties allowed us to limit the parameter scan, removing redundant parameter sets, in the following ways:

- First, we limited the values of $l$ to 0 (no linear contribution) or 1 (linear contribution), and only scanned the values of $c$ and $n$, relative to these fixed linear contributions. Therefore, for all model including a linear component, $l$ = 1, and the values of the other parameters indicate the relative contribution of the linear component.
- Further, for parameter sets where $l$ = 0, we likewise fixed the value of $c$ to 1 and only scanned $n$ to determine the relative contributions of the circadian and noise processes. Thus, for any patient whose pattern of dynamics was categorised as "circadian" (with no linear contribution), $l$ = 0 and $c$ = 1 for the model parameters.

#### *Modelling results*

As a reminder, at each set of parameters, seizure dissimilarities and the corresponding temporal correlation patterns were simulated 1000 times. Each simulated temporal correlation pattern was then compared to the patient's observed temporal correlation pattern by computing the mean squared error (MSE) between the simulated and observed patterns. The likelihood $L$ of a given parameter set was defined as the percentage of "good matches" (simulations with MSE ≤ 0.02185) produced by the 1000 noisy simulations at those parameter values. A model was termed a "linear model" if $c$ = 0, a "circadian model" if $l$ = 0, and a "linear + circadian model" if $l$ > 0 and $c$ > 0. For each class of model (linear, circadian, or linear + circadian), the model's likelihood ($L_l$, $L_c$, or $L_{l+c}$, respectively) was the highest likelihood among the set of qualifying parameter sets, and the "best model" was the model with the highest likelihood, $L_{max}$. $L_n$ was also defined as the highest likelihood of the parameter sets without any linear or circadian contributions ($l = 0$, $c = 0$, $n > 0$).

However, this "best model" was only the "selected" model for the patient if

1) The best model clearly outperformed noise alone ($L_{max} \geq 2L_n$); otherwise, the patient's dynamics were classified as other/indeterminate.
2) The performance of the linear model and circadian model were clearly distinguishable ($L_l \geq 2L_c$ if the linear model was best; $L_c \geq 2L_l$ if the circadian model was best); otherwise, the patient's dynamics were classified as other/indeterminate.
3) If the best model was linear + circadian, it clearly outperformed the two simpler models ($L_{l+c} \geq 2L_l$ and $L_{l+c} \geq 2L_c$); otherwise, the patient's dynamics were classified as the simpler model (if one simpler model performed comparably by this criterion) or as other/indeterminate (if both simpler models performed comparably).

Fig. S8.1 shows the final modelling results for each patient after these model selection criteria were applied. From the simulated temporal correlation patterns (Fig. S8.1B), it is apparent that our simple model can approximately reproduce the observed temporal correlation patterns (Fig. S8.1A). Fig. S8.1C provides the selected model parameters for each patient, which were chosen using the above criteria. Fig. S8.1D provides the likelihood of the selected model, and Fig. S8.1E shows the relative performances of the "best model," with $L_{max}$, compared to each of the model categories. The relative performance was computed as the likelihood of the best-performing model, $L_{max}$, divided by the likelihood of the given model category of model (see Fig. S8.1 caption for an example). Lower values indicate better performances compared to the best model; indeed, the relative performance of the best model is 1 because $L_{max}/L_{max} = 1$. These relative performances were used to select the final model parameters (shown in Fig. S8.1C) and corresponding model category. For example, in some patients (e.g., Study 012-2, second row from top), the linear + circadian model performed best ($L_{max} = L_{l+c}$), but the linear model was selected because it performed comparably: in this case, $L_{l+c}/L_l = 1.058$, indicating that almost as many simulations from the linear model provided a good match to the observed temporal correlation patterns.

To illustrate how the model likelihood was computed, Fig. S8.2 shows the 1000 simulations arising from the selected parameter sets of three patients, 931, 1005, and 1211, which were modelled using the linear, linear + circadian, and circadian models, respectively. The simulations are ordered from lowest to highest MSE and are divided by whether they fall under the MSE threshold for a "good match" to the observed temporal correlation pattern (shown in Fig. S8.1A). Note the similarity of the "good matches" to the observed temporal correlation pattern of each patient. Note that the average MSE of the set of simulations would not necessarily be an appropriate evaluation for model selection because noisy fluctuations can dramatically alter the simulated temporal correlation pattern, especially if the number of analysed seizures was small. For example, for patient 931 (Fig. S8.2A), although some simulations provide a good match to the observed temporal correlation pattern, other simulations have a very high MSE. Our approach determines the "likelihood" of observing such good matches to the observed dynamics, without penalising the parameter set for also producing drastically different dynamics under different noise realisations.

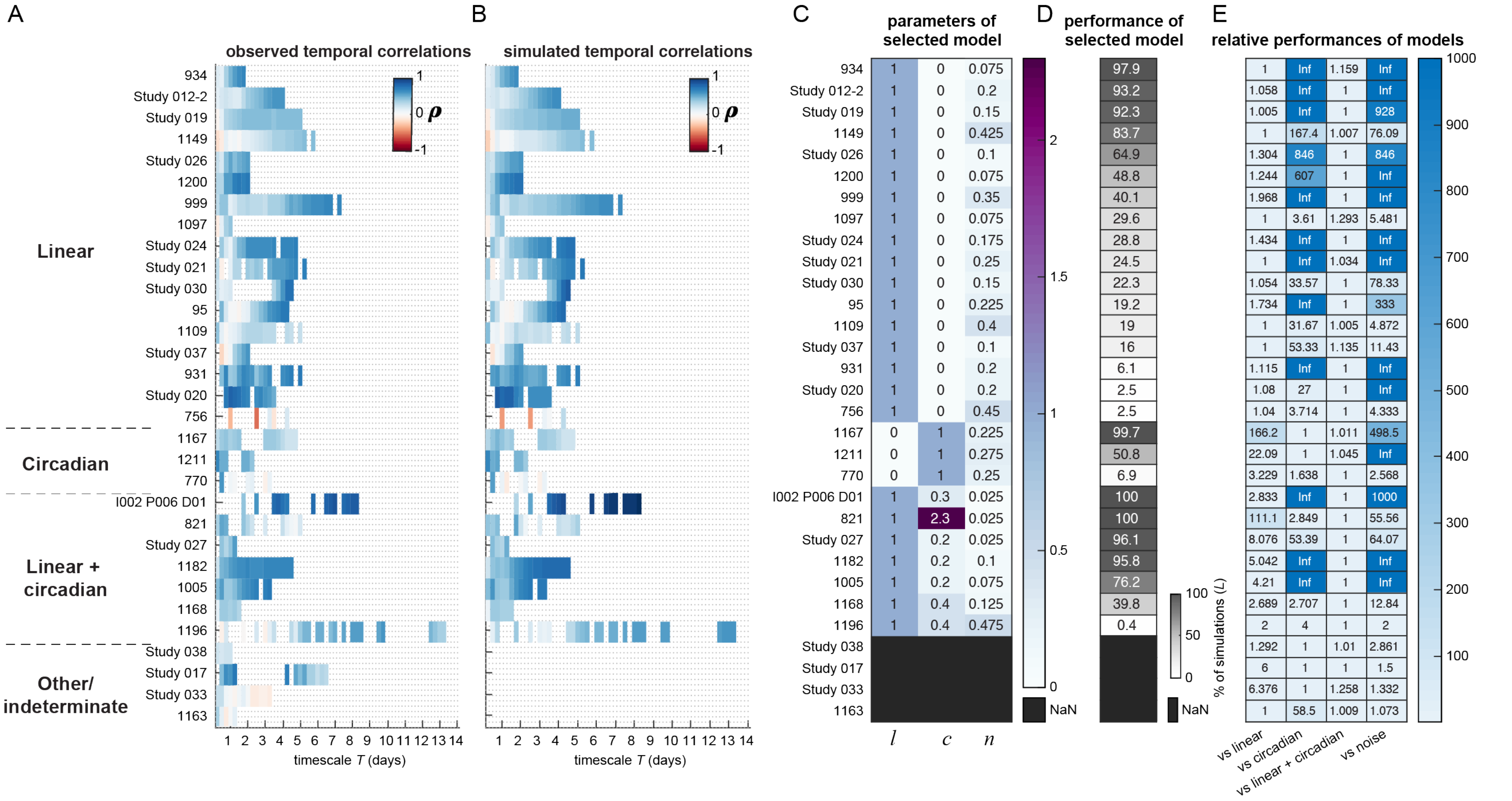
A
B
C
D
E
observed temporal correlations
simulated temporal correlations
parameters of selected model
performance of selected model
relative performances of models
Linear
Circadian
Linear + circadian
Other/ indeterminate
934 1 0 0.075 97.9 1 Inf 1.159 Inf
Study 012-2 1 0 0.2 93.2 1.058 Inf 1 Inf
Study 019 1 0 0.15 92.3 1.005 Inf 1 928
1149 1 0 0.425 83.7 1 167.4 1.007 76.09
Study 026 1 0 0.1 64.9 1.304 846 1 846
1200 1 0 0.075 48.8 1.244 607 1 Inf
999 1 0 0.35 40.1 1.968 Inf 1 Inf
1097 1 0 0.075 29.6 1 3.61 1.293 5.481
Study 024 1 0 0.175 28.8 1.434 Inf 1 Inf
Study 021 1 0 0.25 24.5 1 Inf 1.034 Inf
Study 030 1 0 0.15 22.3 1.054 33.57 1 78.33
95 1 0 0.225 19.2 1.734 Inf 1 333
1109 1 0 0.4 19 1 31.67 1.005 4.872
Study 037 1 0 0.1 16 1 53.33 1.135 11.43
931 1 0 0.2 6.1 1.115 Inf 1 Inf
Study 020 1 0 0.2 2.5 1.08 27 1 Inf
756 1 0 0.45 2.5 1.04 3.714 1 4.333
1167 0 1 0.225 99.7 166.2 1 1.011 498.5
1211 0 1 0.275 50.8 22.09 1 1.045 Inf
770 0 1 0.25 6.9 3.229 1.638 1 2.568
I002 P006 D01 1 0.3 0.025 100 2.833 Inf 1 1000
821 1 2.3 0.025 100 111.1 2.849 1 55.56
Study 027 1 0.2 0.025 96.1 8.076 53.39 1 64.07
1182 1 0.2 0.1 95.8 5.042 Inf 1 Inf
1005 1 0.2 0.075 76.2 4.21 Inf 1 Inf
1168 1 0.4 0.125 39.8 2.689 2.707 1 12.84
1196 1 0.4 0.475 0.4 2 4 1 2
Study 038 1.292 1 1.01 2.861
Study 017 6 1 1 1.5
Study 033 6.376 1 1.258 1.332
1163 1 58.5 1.009 1.073
1 2 3 4 5 6 7 8 9 10 11 12 13 14
timescale T (days)
1 0 -1 ρ
l c n
2 1.5 1 0.5 0 NaN
100 50 0 % of simulations (L) NaN
vs linear
vs circadian
vs linear + circadian
vs noise
1000 900 800 700 600 500 400 300 200 100

**Fig. S8.1: Modelling results for each patient.** A) Observed temporal correlation patterns of all patients, sorted by the selected model (linear, circadian, linear + circadian, or other/indeterminate). Note that these results are also included in Fig. 4C of the main text. B) For each patient, the simulated temporal correlation pattern, arising from each patient's selected model parameters (see Fig. S8.1 C), that best matched the observed temporal correlation pattern (i.e., the pattern that had the lowest MSE across the 1000 simulations produced by the selected parameter values). No simulated pattern is shown for the last four patients because none of the models (linear, circadian, or linear + circadian) met the model selection criteria. C) Parameter values of the selected model for each patient, with $l$ controlling the linear contribution, $c$ controlling the circadian contribution, and $n$ controlling the amount of noise added to the simulated changes in dynamics. D) The performance of the selected model, as defined by the likelihood $L$ of the selected model. The likelihood L is the percentage of simulations, created from different noise realisations using the selected model parameter values, that have a MSE $\leq$ 0.02185 to the patient's temporal correlation pattern. A higher likelihood indicates that the model is more likely to produce the patient's observed temporal correlation pattern. E) The relative performances of the different model categories (linear, circadian, and linear + circadian, as well as the noise-only model) compared to the best-performing model. The relative performance is defined as the likelihood of the best-performing model, $L_{max}$, divided by the likelihood of the given type of model. Lower values indicate better performances compared to the best model. For example, for patient 934 (top row), the linear model performed best ($L_{max}/L_l = 1$ because $L_{max} = L_l$). None of the simulations arising from the circadian or noise models provided good matches to the observed temporal correlation pattern, so the relative performance of these models is positive infinity ($L_{max}/0 = +\infty$). Finally, the linear model performed 1.159 times better than the linear + circadian model. Although the likelihood of these two models was therefore similar, the simpler model (with $c$ = 0) provided a more parsimonious explanation of the dynamics. For each patient, the relative model performances were used to determine the model that clearly outperformed the other categories of models, while also providing the most parsimonious explanation of the observed dynamics.

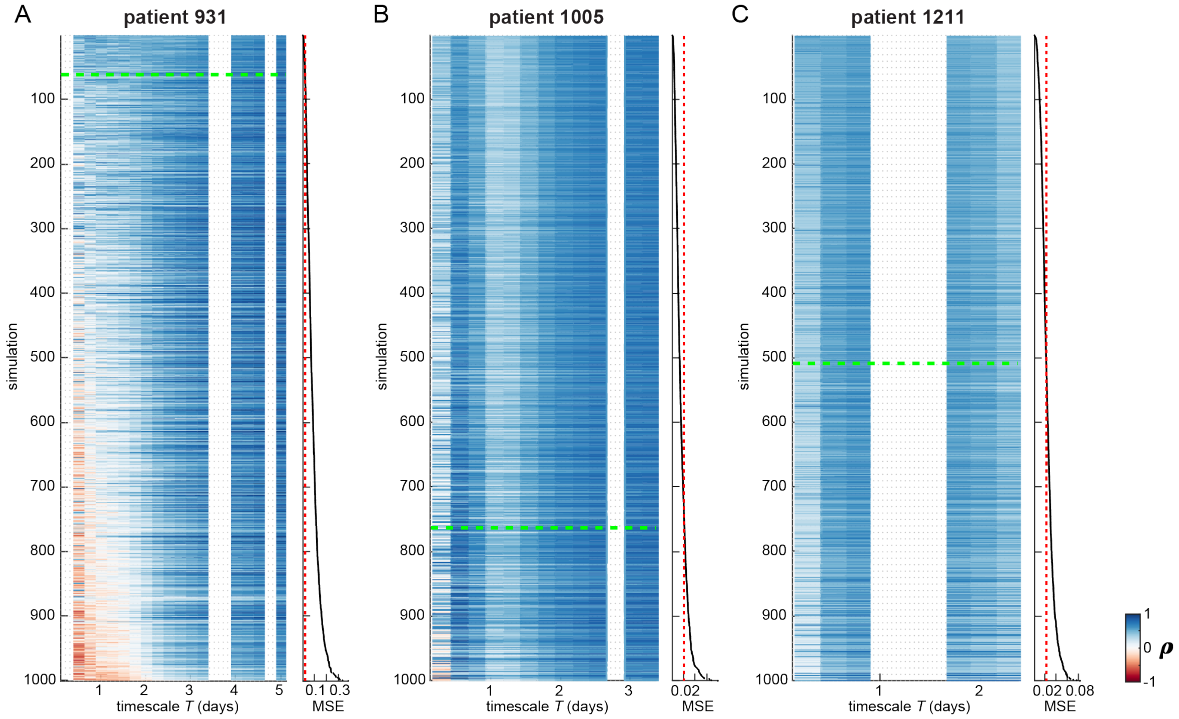


**Fig. S8.2: All simulated temporal correlation patterns arising from the selected model parameters of three example patients: A) patient 931, B) patient 1005, C) patient 1211.** See Fig. S8.1C for the parameters used to generate the simulated temporal correlation patterns. From left to right, the example patients were categorised as linear, linear + circadian, and circadian. For each patient, the heatmap shows the temporal correlation patterns, with the MSE of each temporal correlation pattern (compared to the patient's observed temporal correlation pattern) to the right of the heatmap. Simulations are ordered from lowest MSE (top) to highest MSE (bottom). The red dotted line shows the MSE threshold for a "good match" to the observed temporal correlation pattern, and the dotted green line marks the boundary of the simulations that fall under this threshold.

**S9: Hypothesised model for generating variability in seizure pathways**

Fig. S9 shows a model for how variability in within-patient seizure pathways could arise from a limited number of potential seizure network dynamics. The hypothesised model also accounts for why elements of seizure dynamics are conserved across subsets of seizures. For simplicity, seizure network dynamics are represented as network states, each of which represents a different pattern of functional interactions between the recorded brain areas. The order of network states can be then considered a state progression that defines the seizure pathways.

In the proposed model, various decision points, existing on the framework of potential seizure pathways, produce a repertoire of seizure state progressions. While some parts of the progressions appear deterministic (one network state always leads to a certain subsequent state), at other times a decision point may determine 1) the seizure onset state, 2) the next network state, if multiple progressions are possible, or 3) whether the seizure terminates early in the state progression.

This model would also explain why seizure variability can either manifest as relatively distinct seizure types or as a spectrum of dynamics in our data. A greater number of decision points, which in turn produce a range of small variations between seizures, would produce a spectrum of seizure dynamics. Fewer decision points and/or separate seizure pathways could produce groups of seizures that each have a characteristic state progression.

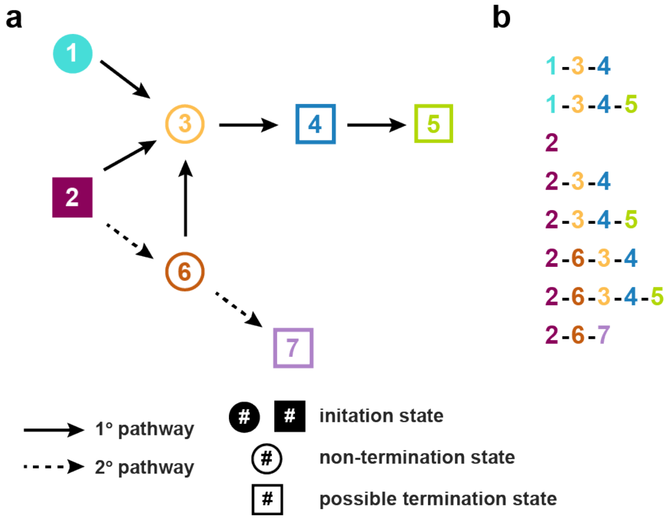


**Fig. S9. Hypothesised model for generating variability in seizure pathways.** (a) Diagram of possible seizure pathways, which are described as transitions between seven network states. For simplicity, we use a schematic of seizure evolution that provides examples of seizure variability features observed in our data. States that are filled in (states 1 and 2) are possible initiation states in the seizure pathway. Dotted arrows represent secondary transitions that are less likely to occur. Square states indicate points in the progression where the seizure may terminate. While some transitions are deterministic (e.g., state 3 always progresses to state 4), other states are decisions points at which variability is introduced into the seizure progression. Variability can be introduced by alternative onsets (e.g., onset states 1 and 2, which can both lead to state 3), different possible progressions (e.g., state 6 can progress to either state 3 or 7), and potential termination points (e.g., state 4 can terminate the seizure or progress to state 5). (b) Potential seizures arising from these seizure pathways, demonstrating variability in state onset, state progression, state termination, and state inclusion. All these types of variability are observed in our cohort. Note that the last three progressions, beginning with the state sequence (2, 6), will be rarer since these transitions are less likely.

**Text S10: Dimensionality reduction using non-negative matrix factorization**

Non-negative matrix factorisation (NMF) was used to reduce noise in each patient's connectivity matrix, $V$, in which each column corresponded to the functional connectivity of a seizure time window. NMF factored $V$ into two non-negative matrices, $W$ and $H$, such that $V \approx W \times H$. The matrix $W$ contained patient-specific basis vectors, each of which had $6*(n^2 - n)/2$ features that captured a pattern of connectivity across all channels and frequency bands. Each original ictal time window was summarised as an additive combination of these basis vectors, with the coefficients matrix $H$ giving the contribution of each basis vector to each time window.

To determine the optimal number of basis vectors, $r$, for each patient, the highest $r$ that produced consistent sets of basis vectors was found (Fig. S10.1). This approach, known as stability NMF (1), exploits the non-deterministic nature of NMF to identify the $r$ at which $W$ consistently converges to a similar set of basis vectors. Since the resulting stable NMF basis vectors can be reliably found, they are thought to provide a meaningful representation of the data. To perform stability NMF for each patient, the value of $r$ was scanned from 1 to 20. This scan range was chosen based on the observation that the stability of the factorisation greatly decreases at approximately $r>10$ in our data, and is consistent with the number of connectivity patterns typically found in ictal iEEG data in other studies (2–4). At each $r$, NMF of $V$ was performed 25 times using the alternating nonnegative least squares with block principal pivoting method (5, 6). Each iteration used different random initializations of $W$ and $H$, thus yielding 25 different factorizations of $V$ at each value of $r$. Using the method established by Wu *et al.* (1), for each $r$, the instability $I$ of two sets of basis vectors $W$ and $W'$ was defined as

$$I(r)_{W,W'} = \frac{1}{2r}\left(2r - \sum_{j=1}^{r} \max_{1 \le i \le r} P_{ij} - \sum_{i=1}^{r} \max_{1 \le j \le r} P_{ij}\right)$$

where $P$ is the Pearson's cross-correlation matrix of the sets of basis vectors. Low values of $I$ indicate that similar sets of basis vectors were found in the separate iterations; indeed, if the two sets of basis vectors are the same (minus reordering), then $I = 0$. The instability of all 25*(25-1)/2 pairs of basis vector sets was then averaged to produce $I_{avg}(r)$. The highest $r$ with $I_{avg}(r) \le 0.005$ was selected for each patient, thus allowing small deviations between the observed basis vector sets, while still enforcing consistent factorisations across iterations. At this $r$, the factorisation yielding the lowest reconstruction error was used to construct $V^* = W \times H$, a lower-rank approximation of the original time-varying seizure functional connectivity. This noise-reduced version of the connectivity was used in the downstream analysis.

Note that the NMF factorisation can also be used to cluster seizure time windows into states (7, 8) (Fig. S10.2), and we use this approach to provide an alternative visualisation of seizure pathways on Zenodo (http://dx.doi.org/10.5281/zenodo.3560736).

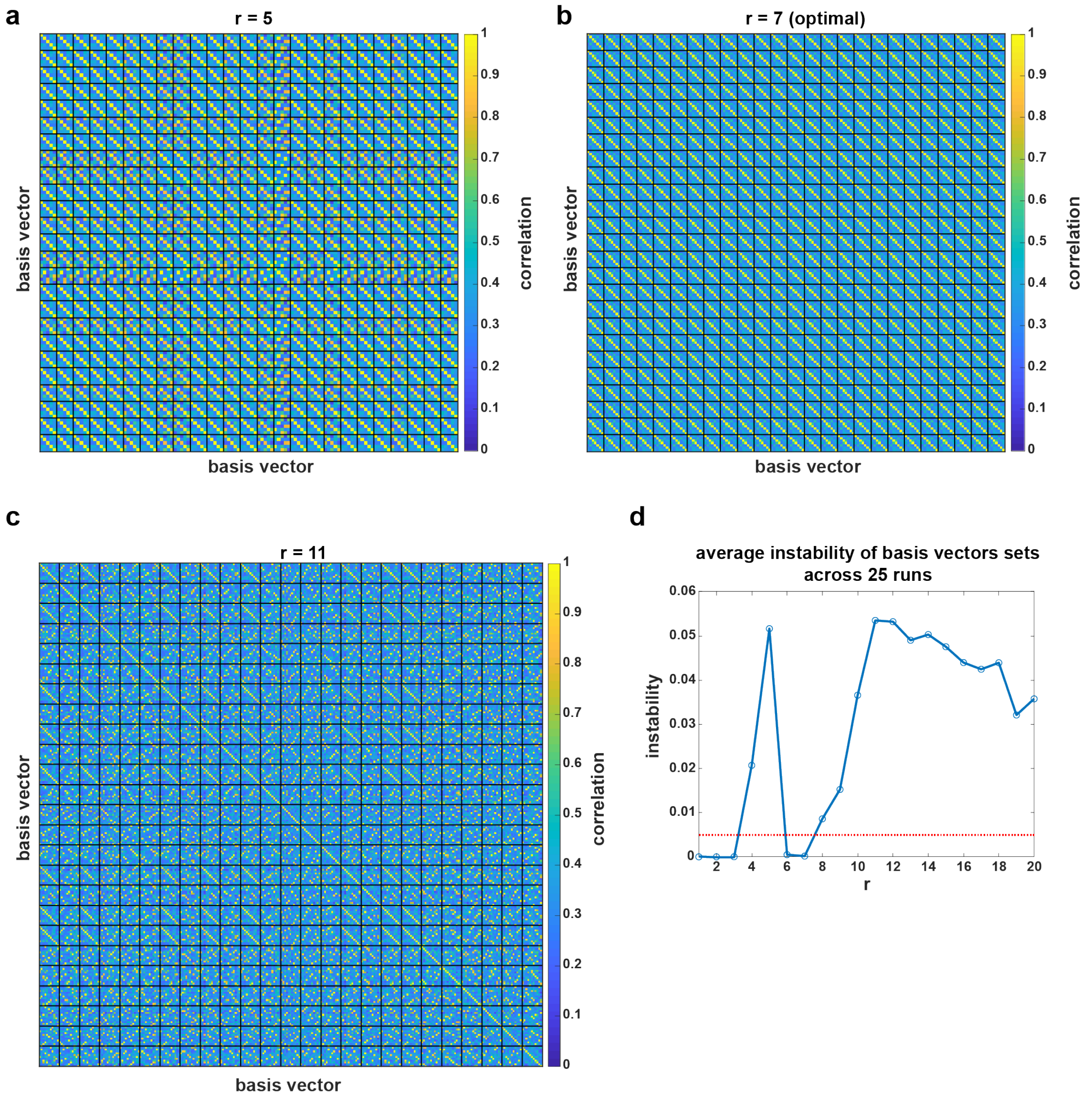


**Fig. S10.1: Finding the optimal number of NMF basis vectors using stability NMF in an example patient, I002_P006_D01.** The optimal number of states was 7 in this patient. At each number of basis vectors, NMF was repeated 25 times using different initialisations, yielding 25 sets of basis vectors. (a-c) Correlation matrices showing Pearson's correlation between all pairs of basis vectors found within and between different initialisations of the NMF algorithm. In each matrix, the same number of basis vectors were optimised for in each run: 5 (a), 7 (b), or 11 (c) basis vectors. Black lines mark the divisions between different runs of the algorithm. When possible, the basis vectors were re-ordered to emphasise similarity between different runs. Specifically, within run *i*, if there was a unique closest match to each of the basis vectors of run 1, the basis vectors of run *i* were re-ordered so that they were in the same order as the corresponding basis vectors in run 1. When r = 7 (b), note that similar sets of basis vectors were found across each run: in a given run, each basis vector had a high (close to 1) correlation to a basis vector of another run. Meanwhile, when r = 5 (a) or r = 11 (c), there was variability in the sets of basis vectors found across runs. (d) Plot of the instability, I, of the basis vector sets, averaged across all pairs of runs, vs. the number of basis vectors, r. For a given r, the average instability quantifies the dissimilarity in the basis vectors found across runs. A low average instability indicates that similar sets of basis vectors were found regardless of the initialization of NMF algorithm; i.e., NMF converged to similar sets of basis vectors from different initial points in the search space. We defined the optimal number of basis vectors, r, as the highest r at which $I(r) < 0.005$. For this patient, the optimal number of basis vectors was $r = 7$.

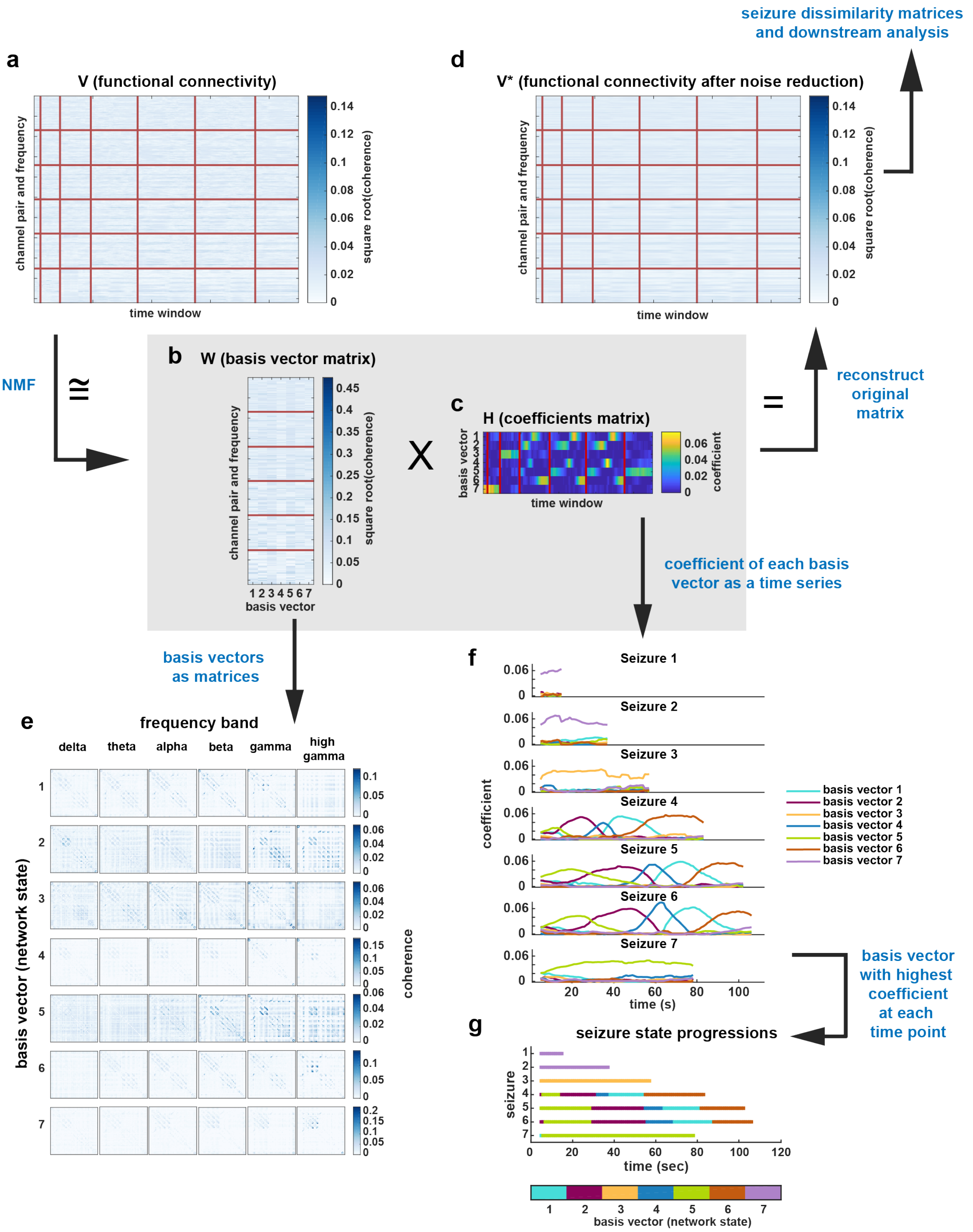


**Fig. S10.2: Workflow for using NMF to reconstruct seizure functional connectivity and assign network states to each time point (alternative visualisation of seizure pathways).** The factorisation of patient I002_P006_D01, which was chosen using stability NMF (see Fig. S10.1), is used as an example. From the matrix V (a), which contains the functional connectivity evolution of all seizures, NMF found a set of basis vectors, each of which forms a column in the matrix W (b), and a coefficients matrix, H (c), that describes the contribution of each basis vector to the observed connectivity at each time point. W and H were used to construct a noise-reduced version of the original matrix, which we call V* (d). In V, V*, and W, rows correspond to features (here, the coherence between a given pair of channels at a given frequency band). The boundaries between features corresponding to different frequency bands are delineated using horizontal red lines (e.g., the top rows correspond to coherence in the delta frequency band). Note that the heatmap colouring corresponds to the square root of the coherence between pairs of channels to aid in visualising the structure of the data; however, the original coherence values were used for NMF and the downstream analysis. Meanwhile, the columns of V, V*, and H correspond to a seizure time window, with vertical red lines marking the boundaries between seizures. (e) Each NMF basis vector is a single column vector in W; however, each of the component frequency band matrices can be re-written in matrix form to emphasise the network pattern. Each basis vector therefore corresponds to a set of six connectivity matrices that describe the network configuration of all pairs of channels across the six frequency bands. Note the different colourbar scale for each basis vector. (f) The coefficients matrix H can be visualised as a time series of the coefficients for each basis vector. Each time point corresponds to the functional connectivity of a 10 s window of ictal iEEG and, at each time point, the coefficients indicate how much each basis vector contributes to the observed seizure connectivity. Note that at a given time, a single NMF basis vector usually has a much higher coefficient relative to the other basis vectors (i.e., there is sparsity in each column of the coefficients matrix, H). Thus, the dominant basis vector provides a simplified description of the network dynamics at that time point. (g) As an alternative visualisation of seizure pathways, each time point was assigned to a network state corresponding to the dominant NMF basis vector, resulting in a series of state progressions for each seizure.

**S11: Comparison of seizure dissimilarity to metric distances**

***Seizure dissimilarity is a nonmetric measure***

Our seizure dissimilarity measure is computed by using dynamic time warping (DTW) to align similar sections of seizure dynamics in a pair of seizures, and then taking the average distance between the warped time courses (see main text Methods and Supplementary S2). We wish to point out as a technical note that due to the warping step, the seizure dissimilarity measure is not a metric distance. Like a metric distance, all dissimilarities are non-negative, the dissimilarity of a seizure to itself is zero, and the dissimilarity between pairs of seizures is symmetric; however, the triangle inequality does not necessarily hold. In particular, any two seizures that follow approximately the same pathway will have a near-zero dissimilarity, regardless of their rates of progression along the pathway. However, their relationship to other seizures that share *part* of the same pathway will depend on how long (temporally) the seizures share the same pathway. Thus, although pairs of seizures may have a low dissimilarity, their relationships to other seizures may differ due to their different rates of progression. These situations can, in turn, lead to violations of the triangle inequality.

To illustrate this point, consider two seizures, A and B, that follow the same dynamical pathway, but at different rates; they will be considered virtually equivalent, and thus have a low seizure dissimilarity, due to the dynamic time warping step. However, their dissimilarity to a third seizure, C, that shares only *part* of the same pathway will differ, as their different rates of progression result in different relative durations of disparate dynamics in these comparisons. In such situations, violations of the triangle inequality may arise. These points are illustrated in the hypothetical dissimilarity matrix, D:

| | A | B | C |
|---|---|---|---|
| A | 0 | 0.5 | 3 |
| B | 0.5 | 0 | 5 |
| C | 3 | 5 | 0 |

Seizures A and B are very similar ($d_{AB}$ = 0.5), but seizure A is less dissimilar (i.e., more similar) to C than seizure B is to C, a situation that can arise if seizures A and B progress at different rates. As such, $d_{AB} + d_{AC} = 0.5 + 3 = 3.5$, which is less than $d_{BC} = 5$, violating the triangle inequality.

***Comparison to metric distances***

To evaluate whether the nonmetric nature of seizure dissimilarities affects our analysis, we also compared the within-patient seizures of our cohort using two distance measures of curves/trajectories that are independent of the time series duration as well as the distance traversed by the trajectory in the feature space: the Fréchet distance and the Hausdorff distance. For clarity, we will refer to our original seizure dissimilarity measure as "DTW dissimilarities" in the remainder of this section.

The first alternative dissimilarity measure is the "Hausdorff distance". To compute this measure, the distances between all pairs of time points in the two time series are first calculated. Then, for every time point, the smallest distance to the other trajectory is found. The Hausdorff measure is then the maximum of all of these smallest distances.

Formally, the Hausdorff distance $d_H$ is defined for two curves X and Y in a metric space (M,d) as:

$$d_{\mathrm{H}}(X,Y) = \max\{\,\sup_{x\in X}\inf_{y\in Y} d(x,y),\ \sup_{y\in Y}\inf_{x\in X} d(x,y)\,\}$$

Note that the Hausdorff distance is a metric distance and is independent of the trajectory length. It is also often applied generally to compare sets (i.e. not just trajectories).

Our second alternative dissimilarity measure is the Fréchet distance. The Fréchet distance is the minimum distance required to link a point travelling along one trajectory to a point travelling along another trajectory. This distance can be conceptualised as the smallest leash length required to walk a dog, given that the dog walker and dog respectively follow the two trajectories of interest. Unlike the Hausdorff distance, but like our DTW-based dissimilarity measure, the Fréchet distance respects the temporal order of progression along the trajectories. However, warping between the two trajectories is allowed in order to minimise the distance between them; i.e., the rate of travel along the trajectories is controlled in order to minimise the distance required to link them. In other words, part of the trajectory can be stretched, but entire sections cannot be repeated; in the mapping from the original trajectory to the warped trajectory, the trajectory indices must monotonically increase.

Formally, the discrete Fréchet distance(18) $d_F$ is defined for two curves X and Y in a metric space (M,d) as:

$$d_F(X,Y) = \inf_{\alpha,\beta}\max_{t\in[0,1]}\{d(X(\alpha(t)), Y(\beta(t)))\}$$

Here, $\alpha$ and $\beta$ are monotonic functions that map $t\in[0\ 1]$ to the start and end of the trajectories X and Y, respectively. The Fréchet distance, again, is a metric distance independent of the temporal durations of and distance traversed by the two trajectories.

Fig. S11A-C compares the three measures of seizure dissimilarity (DTW dissimilarity, Hausdorff distance, and Frechét distance) in our example patient from the main text, patient 931. The agreement between the different measures is visually apparent: all three dissimilarity/distance matrices have similar structures (Fig. S11A-B). There is also a high Spearman's correlation between DTW dissimilarities and each of the other measures (Fig. S11C), indicating that the relative orders of the seizure dissimilarities is mostly retained using the alternative measures. Notably, DTW dissimilarities tend to be smaller than Hausdorff and Frechét distances. This difference is unsurprising given that the Hausdorff and Frechét distances are both highly dependent on the largest distances between the two trajectories, while the DTW measure averages the distances across all warped timepoints. Thus, the DTW measure mitigates the effect of brief, large distances if the trajectories are otherwise similar. Across all patients, DTW dissimilarities also tend to have a high correlation with Hausdorff and Frechét distances (Fig. S11D), indicating that it captures similar information to these metric distances.

To explore the effects of the alternative measures on our main analysis, in each patient we repeated the comparison of temporal distances and seizure dissimilarities with the Hausdorff and Frechét distances (see main text Methods, "Comparison to temporal distances"). For each patient, this analysis resulted in a "temporal correlation" (correlation between temporal distances and seizure dissimilarities) for each measure. Fig. S11E shows scatter plots of the original temporal correlations vs. the temporal correlations computed using the alternative measures. Notably, the temporal correlations are relatively similar regardless of the dissimilarity measure; patients who originally had a positive temporal correlation still show the same relationship using the new measures. Thus, the observed temporal relationships between within-patient seizures are still evident if a metric distance is used to compute seizure dissimilarities. As such, although our nonmetric DTW seizure dissimilarity measure must be used carefully as a substitute for a distance measure, we still find qualitatively similar results if we replace it with a metric distance.

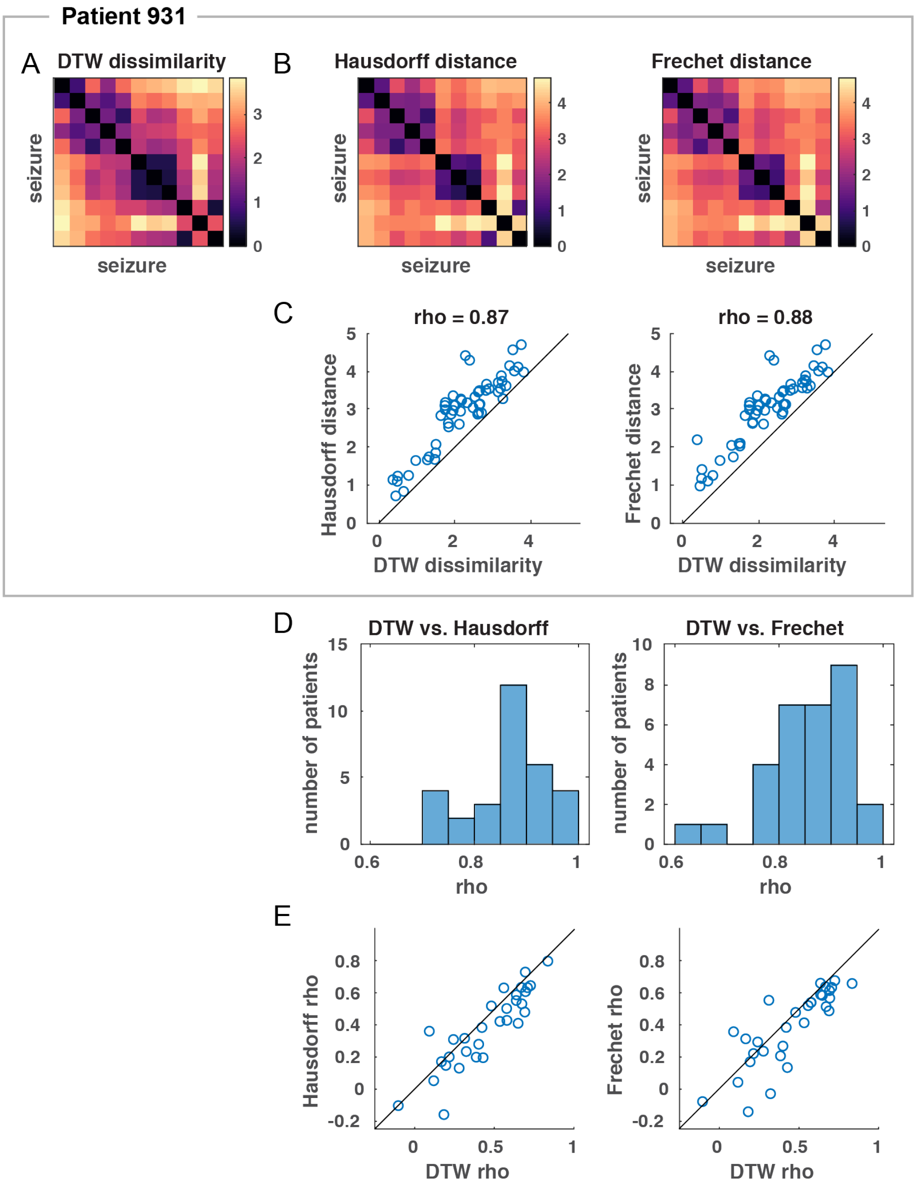


**Fig. S11: Comparison of DTW dissimilarity measure to alternative dissimilarity measures.** A) Seizure dissimilarities of patient 931, computed using our DTW measure of seizure dissimilarity. B) Seizure dissimilarities of patient 931, computed using two alternative measures: the Hausdorff distance (left) and Frechét distance (right). C) Scatter plots of the DTW seizure dissimilarities vs. the Hausdorff distances (left) and Frechét distances (right) of all seizure pairs in patient 931. The black line indicates the equivalence line. Spearman's correlation between each pair of measures is shown above each scatter plot. D) For each patient, Spearman's correlation was computed between the DTW dissimilarities and each of alternative measures for all pairs of seizures. These histograms show the distributions of those correlations across all patients. E) The correlation between each dissimilarity measure and temporal distances ("temporal correlation") was computed for each patient. These scatter plots show the temporal correlation from the DTW dissimilarity measure vs. the temporal correlations computed using each alternative measure of seizure dissimilarity. The black line indicates the equivalence line.
